# Convergent genomic responses of human gut bacteria to variations in industrialization

**DOI:** 10.1101/2025.10.20.683395

**Authors:** Malte Rühlemann, Lénárd L. Szánthó, Silvio Waschina, Lucas Moitinho-Silva, Laura K. Mews, Joan F. Camarena, Hannah Jebens, John Costa, Vanessa Juimo, Alain Fezeu, Adwoa Agyei, Mary Y. Afihene, Shadrack O. Asibey, Yaw A. Awuku, Amoako Duah, Yvonne A. Nartey, Fatimah Ibrahim, Yvonne A. L. Lim, Tan M. Pin, Charles Onyekwere, John Rusine, Ivan E. Mwikarago, John Baines, Andre Franke, Gergely J Szöllősi, Ramnik Xavier, Eric J. Alm, Mathieu Groussin, Mathilde Poyet

**Author notes:** These authors contributed equally to this work (Vanessa Juimo, Alain Fezeu, Adwoa Agyei, Mary Y. Afihene, Shadrack O. Asibey, Yaw A. Awuku, Amoako Duah, Yvonne A. Nartey, Fatimah Ibrahim, Yvonne A. L. Lim, Tan M. Pin, Charles Onyekwere, John Rusine, Ivan E. Mwikarago). co-senior authors (Eric J. Alm, Mathieu Groussin, Mathilde Poyet). Corresponding authors: Mathilde Poyet, Mathieu Groussin and Eric Alm.

## Abstract

To what extent gut bacteria respond to the distinct ecological pressures imposed by human lifestyle remains unclear. Here, we investigate how genomic adaptation in gut bacteria differ between industrialized and non-industrialized human populations. We generated a broad collection of isolate genomes spanning diverse host geographies, lifestyles, species, and strains. We first found that compared to MAGs, paired isolate genomes recover more functional elements and signals of horizontal gene transfers (HGTs). Leveraging isolate genomes from multiple species, we find that strains from industrialized hosts experience an expansion of proteome size and harbor greater pangenome fluidity, driven by recent events of HGTs. Gene- and variant-level analyses reveal convergent patterns of lifestyle-specific adaptation in functions that are critical for ecological adaptation, such as stress response, cell envelope remodeling and central metabolism. Our results demonstrate that industrialization imprints evolutionary signatures on gut bacterial genomes, illuminating the effects of rapidly changing environments on human biology.

## Introduction

Human populations living in industrialized societies harbor gut bacterial communities that differ markedly from those in non-industrialized populations ^1–4^. These differences are driven by variation in lifestyle, diet, sanitation and exposure to environmental microbes ^5,6^. As such, lifestyle-associated factors not only reshape the nutrient landscape available to gut bacteria, but also alter the set of microbial interaction partners with which each species coexists and interacts. While shifts in microbiome composition across human populations – particularly in relation to industrialization and subsistence strategies – are now well documented, far less is known about whether and how individual bacterial strains evolve and adapt to these host-associated environments. In contrast to environmental microbes and pathogens ^7^, adaptive genomic signatures in commensal gut bacteria remain poorly characterized, despite evidence for adaptation occurring within individual hosts on timescales of days to months ^8–12^.

Addressing this gap requires deep, high-quality collections of cultured isolates and their genomes from a wide range of human lifestyles and geographic settings. However, existing culture collections remain heavily biased, primarily representing microbiota from individuals living in countries with high human development index (HDI) such as the United States, China, and European nations ^13–20^. In our previous work ^21^, we began to address this imbalance by generating the first Global Microbiome Conservancy (GMbC) isolate genome collection, comprising over 4,000 gut bacterial genomes from host populations spanning a broad range of geographies and lifestyles. Others also expanded the phylogenetic and geographic diversity of target taxa, such as Segatella copri (previously named Prevotella copri) ^22^.

In addition to isolates, metagenome-assembled genomes (MAGs) have become a widely used resource to explore microbial diversity ^23–26^, including from non-industrialized populations for which access to isolate genomes is more challenging. While MAGs have enabled large-scale surveys of phylogenetic diversity, their use for detecting recent adaptation or fine-scale genomic features remains debated ^27,28^. Potential limitations of MAGs include variable completeness, the inability to fully capture within-host population heterogeneity, and the loss of accessory functions – particularly those associated with mobile genetic elements (MGEs) and horizontal gene transfers (HGTs) ^27,29^. These challenges stem from technical and biological factors: binning errors, reliance on single-copy core genes (SCGs) for estimating completeness, and the inherent difficulty of assembling genomes in the presence of high microdiversity and strain-level variation ^30,31^. To robustly evaluate MAG quality, direct comparison of MAGs to taxonomically-paired isolate genomes from the same sample is required. While efforts have been made in this direction ^28^, such comparisons have so far been limited in scale and taxonomic breadth, leaving a critical gap in our ability to assess how well MAGs represent real genomic diversity.

Here, we expand and leverage the GMbC collection of isolate genomes and MAGs to investigate how industrialization shapes the evolutionary trajectories and adaptive potential of gut commensal bacteria. We found that gut bacterial genomes from industrialized hosts show evidence of proteome expansion and elevated pangenome fluidity, both owing to a recent acceleration in HGT-driven gene acquisition. Across multiple species, we uncover parallel signals of genomic adaptation to host industrialization status, including lifestyle-specific gene enrichment, signatures of positive selection, and convergent non-synonymous single nucleotide variants (SNVs) in genes that are functionally relevant for ecological adaptation.

## Results

### Extensive species, strain and geographic diversity in the GMbC isolate genome collection

To build the GMbC isolate genome collection of human gut bacteria, we employed culturing strategies designed to maximize bacterial species and strain diversity (see Methods) (Fig. 1A). Host individuals from the GMbC cohort ^6^ were selected to represent a broad range of lifestyles, geographic backgrounds and microbiome diversity (see Methods) (Fig. 1B & Supp. Table 1). We release a new set of 1,841 high-quality isolate genomes, which we integrate into our previously published set of 4,140 genomes ^21^, resulting in a total of 5,981 high-quality genomes (see Methods) (Fig. 1C). GMbC genomes have a median completeness of 99.2%, contamination of 1.67%, and quality score of 98.3. Using similarity thresholds at 99% and 95%, GMbC isolate genomes were clustered into 1,133 strain- and 434 species-level genome bins (StGBs and SGBs, respectively). Strain-level genome bins represented in multiple hosts were further split by host to ensure that each StGB represented a host-specific strain (see Methods). These isolates were cultured from stool samples of 56 donors across 24 geographic locations in 9 countries (Fig. 1C, Supp. Fig. 1, and Supp. Table 1). Of the 434 SGBs, 403 were obtained from individuals living more non-industrialized lifestyles, and 102 from individuals in more industrialized settings (Fig. 1D). For comparison, only 106 SGBs were recovered in our previous BIO-ML collection, which included only individuals from industrialized populations in the United States ^12^. The median number of GMbC SGBs per donor, geographic location, self-declared ethnicity, and country is 17, 41, 52, and 87, respectively (Fig. 1D & E).

**Figure 1.**
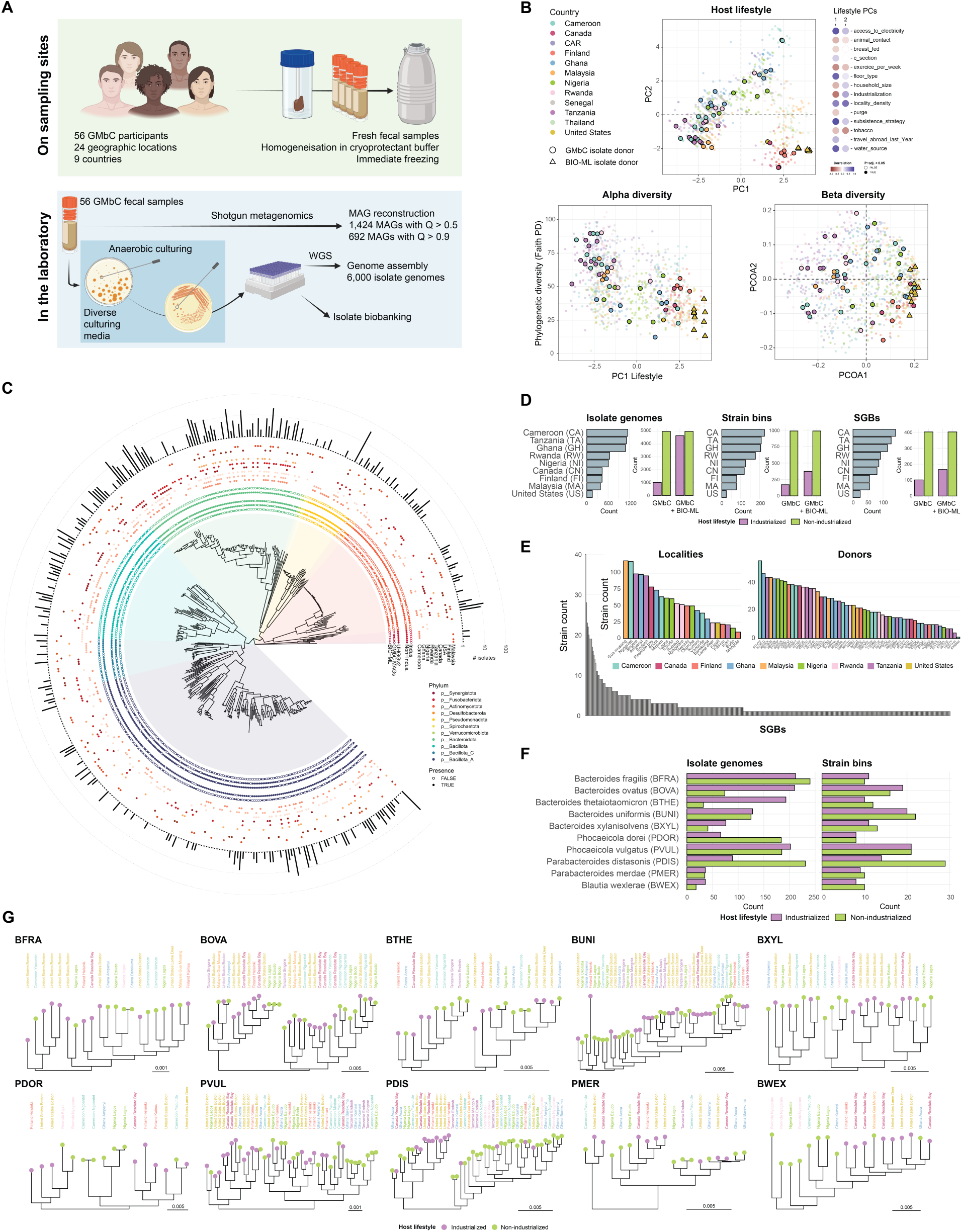
Geographic, species, strain, and host lifestyle diversity in the GMbC collection of human gut bacterial isolate genomes. A. Overview of sampling, preservation, culturing, isolation, and sequencing procedures for gut bacterial genomes (see Methods). B. Lifestyle and microbiome diversity of donors used for culturing and isolating gut bacteria in the context of the broader GMbC + BIO-ML cohort. Top panel: dimensional reduction analysis of various lifestyle factors (see Methods). Donors used for culturing are shown in larger symbols with dark border. GMbC donors are shown in circles, BIO-ML donors are shown in triangles. Spearman correlations between the first two PCs and individual lifestyle factors are shown on the right. Alpha diversity (measured with Faith PD index) and beta diversity (unweighted UniFrac) of GMbC and BIO-ML isolate donors and participants are shown in bottom panels. C. Phylogenomic tree of representative genomes from 434 species-level genome bins (SGBs). Inner ring shows overlap with external genome collections (UHGG v2, GMbC MAGs, BIO-ML). Middle ring indicates host lifestyle origin (industrialized or non-industrialized). Outer ring shows country distribution and isolate genome counts per SGB. Clade colors represent phyla. D. Isolate genome, strain bin, and SGB counts by country and host lifestyle. Strain bins group genomes from the same donor with >99% similarity (see Methods). Counts that include isolate genomes of the BIO-ML collection per host lifestyle are also shown. BIO-ML isolate genomes were generated following the same pipeline as described in A (see Methods). E. Distribution of strain bin counts across SGBs, localities and individual hosts. Colors denote country. F. Ten bacterial species with ≥8 strain bins sampled from industrialized or non-industrialized hosts. Barplots show isolate and strain bin counts per lifestyle. G. Phylogenetic trees of representative strain bin genomes for the 10 species in panel E. Tip points indicate host lifestyle; labels show country/locality and are color-coded by country. Trees are midpoint-rooted. Branch length scales are in expected number of substitutions per site.

Next, we assessed the phylogenetic and functional diversity of the GMbC isolate genome collection. The 434 GMbC SGBs span 13 distinct phyla and 46 families, encompassing both dominant taxa of the human gut microbiome and phylogenetic groups that are rare, underrepresented, or difficult to cultivate (Fig. 1C). Notably, the collection includes isolates from Spirochaetota (*Treponema_D succinifaciens*, n = 5; *Treponema_D peruense*, n = 2) and Synergistota (*Cloacibacillus porcorum*, n = 6). Contrarily to Treponema pallidum, which is a pathogen, very little is known about *Treponema succinifaciens*. It was frequently found to be in relatively high abundance in the gut microbiome of non-industrialized populations, while being absent from those of industrialized populations ^6,32,33^. Treponema in non-industrialized populations is thought to be a commensal that may promote fiber degradation in the context of fiber-rich diets ^1^. Looking at the broader GMbC cohort, we recently found *Treponema succinifaciens* to be negatively associated with intestinal inflammation markers ^6^, and to be negatively associated with hypoxia and TNF signaling gene expression in industrialized contexts ^34^. The *C. porcorum* isolates represent the first cultured strains of human origin, as previously available reference strains (e.g., those in the DSMZ collection) were isolated from pigs ^35^. Furthermore, human-derived *C. porcorum* genomes are absent from large-scale metagenomic resources such as the UHGGv2 MAG collection ^24,36^ (Fig. 1C). The GMbC dataset also includes archaeal isolates, with 7 genomes spanning the phyla Methanobacteriota (Methanobrevibacter_A smithii, n = 2; Methanosphaera stadtmanae, n = 1) and Thermoplasmatota (Methanarcanum hacksteinii, n = 4). Among the 434 SGBs, 55 lack a species-level taxonomic assignment, highlighting gaps in current reference databases such as GTDB (rel214) ^37^. Of these, 27 belong to the Collinsella genus, 2 to Prevotella, and 2 to Faecalibacterium.

The GMbC collection also captures substantial within-species strain genomic diversity (Fig. 1D & E & Supp. Fig. 1). In total, 39 SGBs are represented by at least 10 distinct StGBs, including from multiple key bacterial species in the human gut, such as Bacteroides, Parabacteroides, Blautia species. Genomic-based phenotypic and functional predictions, including amino acid auxotrophies that are associated with chronic diseases ^38^, suggest substantial strain-level diversity among several key taxa, including Prevotella, Veillonella and Blautia species (Supp. Fig. 1). Overall, the GMbC collection extensively samples species and strain-level diversity of human gut bacteria across various human host modalities, including geography and lifestyle, providing unprecedented amounts of genomic material for in-depth genomic and functional investigations of the global gut microbiome. We leverage this phylogenomic diversity in the following analyses to investigate how host industrialization influences the genomic evolution of human gut bacteria.

### Isolate genomes recover more functional and mobile elements than MAGs

We first benchmarked our isolate genomes against metagenome-assembled genomes (MAGs) to assess the advantage of combining MAGs with isolate genomes in our analysis. Previous studies have questioned the completeness and quality of MAGs due to errors introduced during metagenomic assembly and binning, as well as challenges such as within-sample strain diversity ^39^. Although adding MAGs could substantially expand strain representation and species coverage, we wondered whether these hypothesized drawbacks of MAGs could impact our analysis of genomic evolution across host lifestyles, particularly for accessory gene families and mobile genetic elements that may be central to lifestyle-driven adaptive processes. So far, it has remained difficult to systematically evaluate how MAGs compare to isolate genomes because most prior comparisons involved unpaired genomes—i.e., MAGs and isolates obtained from different hosts, and often representing different strains. An advantage of our study design is the ability to directly compare the characteristics of isolate genomes with those of paired MAGs. To do this, we leveraged shotgun metagenomic data that we recently generated from the same fecal samples used to culture our isolates ^6^, and reconstructed MAGs using a multi-binning strategy ^6,40^. We filtered out low-quality MAGs using similar thresholds for completeness (<50%) and contamination (>10%) as for our isolate genomes. This allowed us to assemble a unique dataset of 147 paired MAG–isolate genomes, originating from the same species, and sampled from the same donor (Fig. 2A). These pairs span a broad range of bacterial taxonomies and abundances in the human gut (Fig. 2B & Supp. Fig. 2).

**Figure 2.**
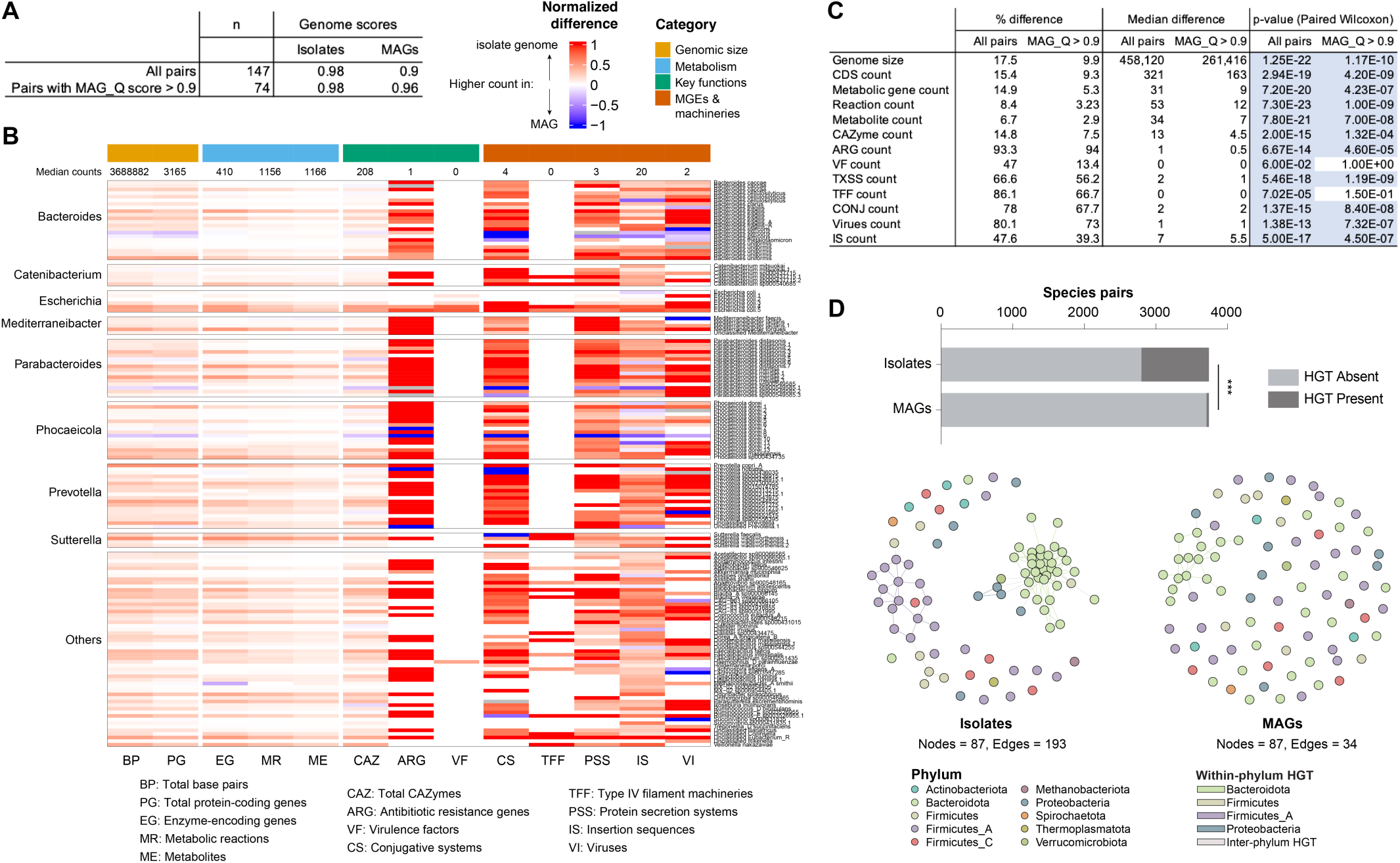
Isolate genomes recover more genomic features and HGT events than MAGs. A. Number and quality scores of MAG–isolate genome pairs. Pairs originate from the same donor sample and species. B. Heatmap comparing genomic feature counts across all genome pairs. Genera and species of genome pairs are shown on the left and right size of the heatmap, respectively. Genomic features are shown in columns, and are grouped in four categories: genomic size, metabolism, key functions and mobile genetic elements (MGEs) & machineries. For each pair, the difference in counts between the isolate genome and the MAG was calculated and normalized to the count in the isolate genome. Features with higher counts in the isolate genome or in the MAG are shown along a gradient of red to blue, respectively. C. Summary statistics of feature differences across all pairs. D. Comparison of HGT events. Between-species horizontal gene transfers (HGTs) were detected across isolate genomes, and across MAGs separately (see Methods). Genomes of MAG-isolate genome pairs cluster in 87 SGBs. Ratio of species pairs with detected HGTs (n >= 1 HGT) were compared with a proportion test (Two proportion Z-test, ***: p = 7.96e-33). Edges in the network indicate that at least 1 HGT was detected between species (nodes).

The median assembly quality score of the 147 MAGs is 0.90, comparable to the UHGGv2 MAG collection (MAG_Q = 0.89) (see Methods). The MAGs show a median completeness of 92.5% and contamination of 2.5% (Supp. Table 2). Among high-quality MAGs (MAG_Q > 0.9; n = 74), the median MAG_Q reaches 0.96 (median completeness = 98% and contamination = 2.5%) – close to that of their paired isolate genomes (Q = 0.98) (Fig. 2A). We used genome-scale metabolic models (GMMs) and reference databases to profile all genomes for functional categories and mobile genetic elements (MGEs) (see Methods).

We found that isolate genomes are significantly larger (paired Wilcoxon test, median difference = 458,120bp, p = 3.6e-22) and contain more coding sequences (CDSs) (median diff. = 321 genes, p = 2.9e-19) than MAGs (Fig. 2B & C, Supp. Fig. 2 & Supp. Table 2). Isolate genomes also harbor more metabolic features predicted by GMMs, including enzyme-encoding genes (median diff. = 31, p = 7.2e-20), reaction pathways (median diff. = 53, p = 7.3e-23), and predicted metabolites (median diff. = 53, p = 7.3e-23) (Fig. 2B & C). This disparity is further pronounced when GMM reconstruction is performed without pathway gap-filling (see Supp. Table 2 and Methods). We also found that MAGs required significantly more pathway gap-filling during GMM reconstruction than isolate genomes (p = 4.7e-12), confirming that gene content inference and metabolic reconstructions are more fragmented in MAGs.

We then compared the counts of specific gene families involved in key bacterial functions, such as carbohydrate metabolism (carbohydrate active enzyme [CAZyme]), antibiotic resistance (antibiotic resistance genes [ARGs]), and virulence (virulence factors [VFs]). These gene families were consistently detected in greater numbers in isolate genomes compared to MAGs (p = 2.0e-15, p = 6.7e-14 and p = 6e-02, respectively) (Fig. 2B & C, Supp. Fig. 2).

We also profiled mobile genetic elements and machinery involved in horizontal gene transfer, including conjugative elements (Conj), insertion sequences (ISs), prophages, type IV filament (TFF) superfamily proteins, and protein secretion systems (TxSS). Each of these categories was more abundant in isolate genomes compared to MAGs (Fig. 2B & C, Supp. Fig. 2).

Importantly, these trends remained consistent even when restricting the analysis to high-quality MAGs (MAG_Q > 0.9; n = 74) (Fig. 2C, Supp. Table 2). Finally, differences in functional content between isolates and MAGs were robust across a range of bacterial taxonomies and relative abundances (Fig. 2B, linear models with taxonomy and abundance, see Supp. Table 2).

Considering that MGEs are better captured in isolate genomes, we next asked whether isolates would also provide greater sensitivity and accuracy for detecting recent horizontal gene transfer (HGT) events. To test this, we applied a previously established BLAST-based pipeline for identifying nearly identical sequences shared between genomes of different species ^21,41^. Our 147 MAG–isolate pairs represent 87 distinct bacterial species. We used this pipeline to detect HGT events across these 87 species using either MAGs or isolate genomes (Methods). Both the number of candidate HGTs (BLAST hits) and the proportion of species pairs involved in putative HGT were significantly higher when using isolate genomes (paired Wilcoxon test, pl = 3.27e-39; Two proportion Z-test, X-squared = 142.4, p = 7.96e-33) (Fig. 2D).

Overall, isolate genomes consistently recover higher numbers of genomic and functional features compared to MAGs. Our results also show that high-quality MAGs still capture much of this information (Fig. 2B & Supp. Table 2). With ultra-deep sequencing, long-read metagenomics and advances in binning algorithms ^2,42,43^, MAGs should achieve performance comparable to isolates in detecting these features. In the following, we chose to consider MAGs for validation analyses: all primary inferences regarding gene content variation and sequence divergence are drawn from the more complete isolate genomes, and MAGs are used when needed to confirm that these patterns persist when sampling a broader set of host individuals (Fig. 7). By leveraging the complementary strengths of both data types, we aim to minimize the risk that assembly artifacts bias our conclusions about lifestyle-associated genomic evolution.

### Bacterial SGBs with broad sampling of StGBs to study the effect of host industrialization status on genomic evolution

To test whether host industrialized vs. non-industrialized lifestyles exert selective pressures that impact genomic evolution and adaptation of gut bacteria, we investigated a range of genomic and phylogenetic features, including gene content, ancestral gene gain and loss events, signals of positive selection, and individual SNVs. We focused on ten bacterial species that were well-represented in the GMbC collection and included a high number of isolate genomes and StGBs from individuals of both lifestyle types (Fig. 1F & G): *Bacteroides fragilis* (BFRA), *Bacteroides ovatus* (BOVA), *Bacteroides thetaiotaomicron* (BTHE), *Bacteroides uniformis* (BUNI), *Bacteroides xylanisolvens* (BXYL), *Phocaeicola dorei* (PDOR), *Phocaeicola vulgatus* (PVUL), *Parabacteroides distasonis* (PDIS), *Parabacteroides merdae* (PMER), and *Blautia A wexlerae* (BWEX). We complemented the GMbC genomes with published isolate genomes of these species from the BIO-ML collection (industrialized host donors from the Boston (MA, USA) area) that we generated using similar culturing and sequencing methods ^12^ (Fig. 1F). For each species, we reconstructed the pangenome (core, accessory and cloud genomes) and the recombination-aware phylogeny of StGB representative genomes (Fig. 1G and Supp. Fig. 3) (see Methods). Phylogenetic signal analyses of the industrialization trait (industrialized / non-industrialized host) using Blomberg’s K revealed that the trait has limited signal in most species (K < 1; Supp. Table 1) and is broadly distributed across StGB phylogenies (Fig. 1G). Only BWEX showed a significant phylogenetic signal (K > 1, p-val < 0.01). Combined with appropriate control for phylogenetic structure in statistical models, these patterns enable the detection of convergent genomic responses to host industrialization status among distantly-related strains, as explored in the following sections.

While we use a binary variable for industrialization status (industrialized vs. non-industrialized host) to investigate patterns of bacterial adaptation, **we acknowledge that this classification oversimplifies a broad spectrum of lifestyle differences**. In particular, non-industrialized populations exhibit greater diversity in subsistence strategies and environmental exposures compared to industrialized populations ^6^. **To address this limitation, we also incorporate a continuous variable derived from a multidimensional reduction of lifestyle-related factors (named “PC1 Lifestyle” thereafter)**, including but not limited to industrialization status (Fig. 1B). This variable was computed from the broader GMbC cohort (n = 1,015 participants) that we recently described ^6^, which includes the GMbC participants used here for bacterial isolation. PC1 Lifestyle is strongly correlated with industrialization status, while also capturing finer-scale variation in lifestyle across participants (Fig. 1B). Where appropriate (i.e. gene enrichment and single nucleotide variant (SNV) analyses in Fig. 5 & 7), we use this higher-resolution variable as a continuous proxy for industrialization to validate associations identified using the binary classification.

### Proteome expansion and increased pangenome fluidity in gut bacteria of hosts with industrialized lifestyles

We first tested whether the total protein-coding gene content differs between strains colonizing individuals from industrialized vs. non-industrialized lifestyles. Across all 10 species examined, strains from industrialized hosts consistently exhibited larger proteome sizes than those from non-industrialized hosts (Fig. 3A). To account for phylogenetic structure, we applied phylogenetic linear regression models across species and identified eight species with significantly larger proteomes in industrialized strains (p < 0.05). Combining species-level p-values using Fisher’s method revealed a strong overall signal of proteome expansion in industrialized populations (p = 6.2e-12) (Fig. 3A).

**Figure 3.**
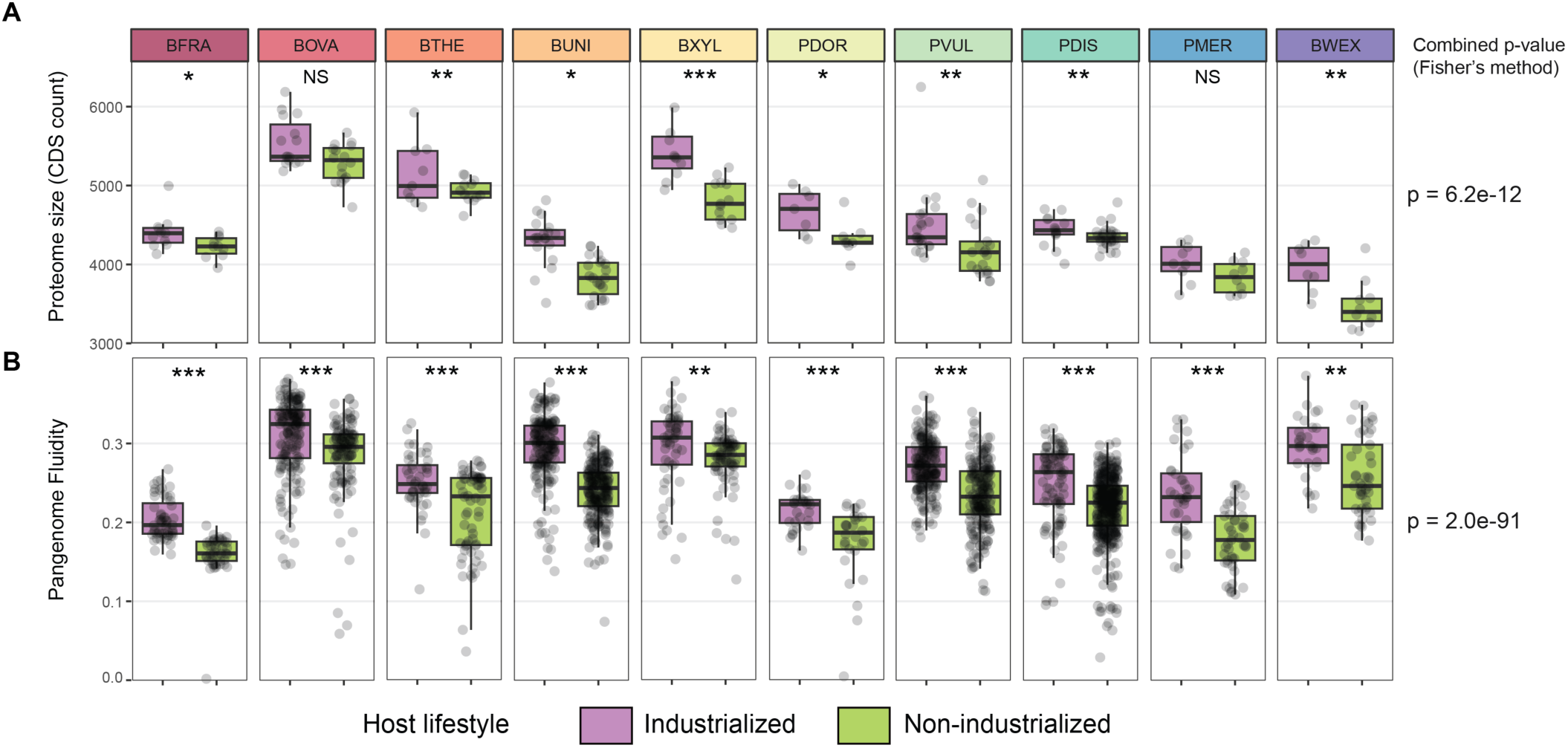
Industrialized host strains exhibit larger proteomes and signatures of relaxed selection. A. Comparison of proteome size (coding gene counts) between strains of host with industrialized vs. non-industrialized lifestyles (in purple and green, respectively) across the 10 species presented in Figure 1. Counts were statistically compared while accounting for phylogeny (phyloglm function, see Methods) (***: p-value < 0.001; **: p-value < 0.01; *: p-value < 0.05; NS: non-significant – this legend applies to all other panels). P-values were combined with the Fisher’s method to test for cross-species evidence of differences in proteome size against the null hypothesis. This p-value is shown on the right of the panel. B. Comparison of pangenome fluidity among industrialization- and non-industrialization-associated strains. The ratio of shared genes was calculated for strain bin pairs, using representative genomes. P-values were combined with the Fisher’s method (p-value shown on the right of the panel)

This proteome expansion may be associated with increases in pangenome size and fluidity. We first confirmed that core genome size does not differ between strains from industrialized and non-industrialized hosts (paired Wilcoxon test, p = 0.92), as expected, indicating that the observed proteome expansion is not driven by changes in conserved genes. We then assessed pangenome fluidity, which quantifies the variability in gene content among strains of the same species. Specifically, fluidity measures the proportion of genes that are not shared between genome pairs, relative to the total number of genes (see Methods). Higher pangenome fluidity reflects a more open and dynamic pangenome, characterized by extensive gene turnover and accessory genome diversity. Across all ten species analyzed, we consistently observed higher pangenome fluidity among strains from industrialized hosts (p-val < 0.01 for each species; Fisher’s combined p-val = 2.0e-91) (Fig. 3B), suggesting that bacterial strains in industrialized environments experience stronger gene turnover and greater genomic plasticity. These findings align with our previous work showing elevated rates of horizontal gene transfer (HGT) in gut bacteria from industrialized populations ^21^.

### Recent increase in rates of bacterial gene gains among industrialized hosts

We next sought to test whether the observed increase in proteome size among strains from industrialized hosts is driven by elevated rates of gene acquisition or by reduced levels of gene loss, relative to strains from non-industrialized hosts. Moreover, it remains unclear whether this expansion reflects a recent evolutionary response to industrialized lifestyles. If the latter were the case, we would expect gene gains and proteome size increases to be confined to terminal branches of bacterial species trees, corresponding to more recent evolutionary events. Alternatively, these events could have occurred along ancestral lineages, with ecological niche selection favoring the colonization of industrialized hosts by strains of larger proteomes.

To distinguish between these scenarios, we used a species tree-gene tree phylogenetic reconciliation method implemented in AleRax ^44^ to identify the branches of the species trees along which genes were gained or lost. AleRax distinguishes two processes of gene gain, either via horizontal transfer or by origination. In order to reconstruct gene trees, we considered all gene families being present in at least 4 different genomes across StGBs. We inferred whether ancestral nodes along the species phylogeny were from industrialized or non-industrialized hosts using Wagner parsimony. We then quantified gene gain/loss events and copy numbers across internal and terminal branches based on reconciliation scenarios, stratified by host lifestyle (See Methods, Supp. Table 4). We found higher counts of gene gains along branches associated with industrialized lifestyles, that gene gains outnumbered losses across most species, and that they mainly occurred along terminal branches (Fig. 4A, Wilcoxon-tests, plain circles show species with statistically significant differences (p-val < 0.05) in event counts between lifestyles). Interestingly, the higher rate of gene gain in *B. thetaiotaomicron* is amplified by high rates of gene gains along ancestral branches associated with industrialized lifestyle, and higher rates of gene loss along ancestral branches associated with non-industrialized lifestyle (Fig. 4A).

**Figure 4.**
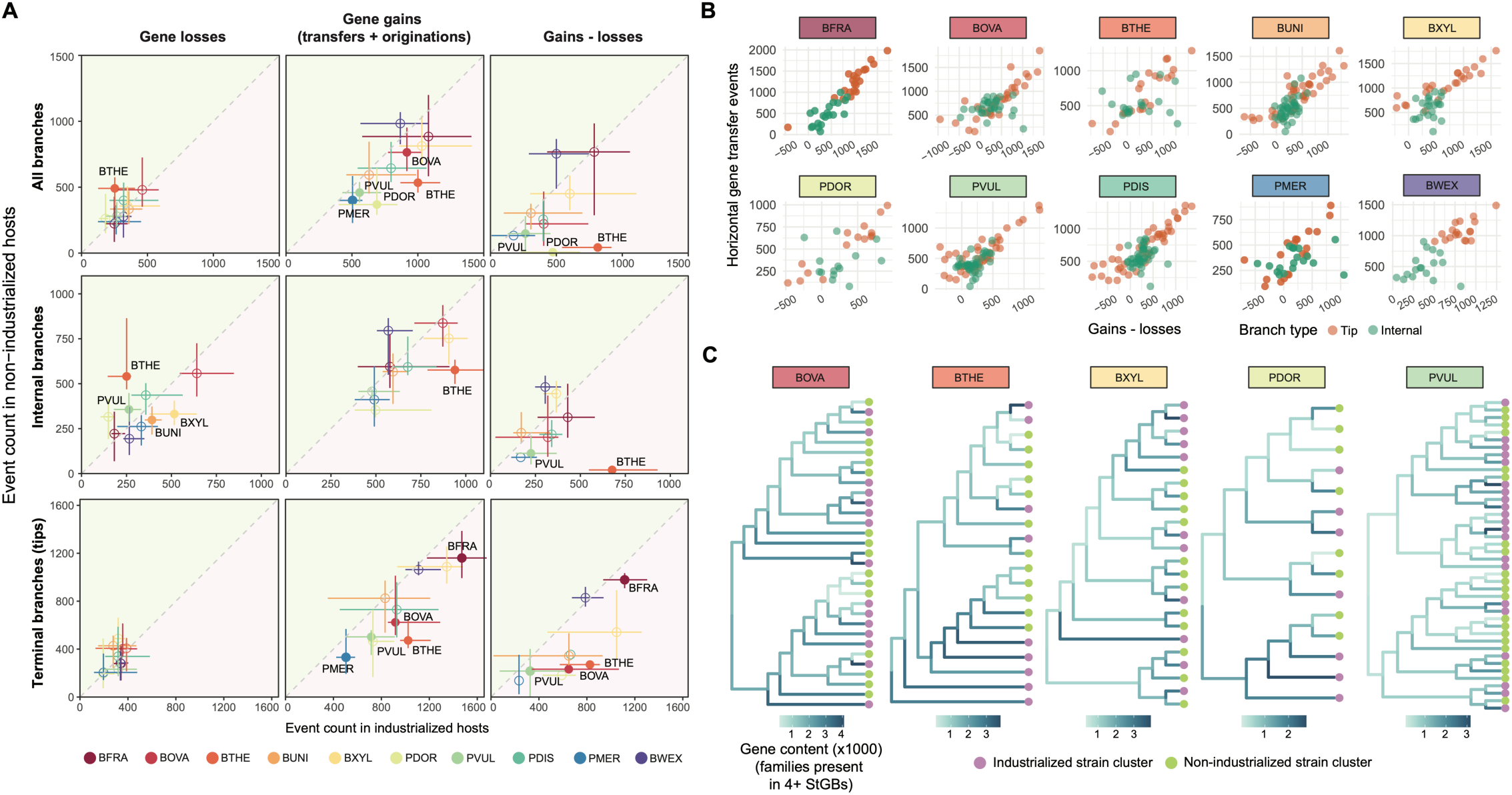
Recent and HGT-driven gene gains promote proteome expansion in industrialized strains. A. Species tree - gene tree reconciliations were sampled to detect and count per-branch events of gene transfer, loss, origination and speciation (see Methods). Counts of per-branch gene loss (left column) and gain (middle), and differences between gain and loss counts (right) were compared between host lifestyle categories (industrialized: purple area; non-industrialized: green area). Gene gains were defined as the sum of gene transfer and origination events. Top row: counts aggregated across all branches. Middle row: Counts of internal branches. Bottom row: counts of terminal (tip) branches. For each species, median counts are shown, with intervals ranging from the 25th to the 75th quantiles. Plain points indicate species for which the difference in counts between host lifestyle categories is significantly different (Wilcoxon tests). Species are colored-coded. B. Correlation between per-branch gain–loss differences and HGT counts, broken down by internal (green) and terminal (orange) branches. All correlations are statistically significant (p-val < 0.001; Spearman correlation tests). C. Increasing gene content along lineages of industrialized hosts. The panel depicts the evolution of the number of genes along the phylogeny of BOVA, BTHE, BXYL, PDOR and PVUL, based on the reconciliation-aware reconstruction of ancestral gene contents. Reconciliations were calculated from the set of gene families present in at least four StGBs (tips of the tree). These 5 species have significant differences in proteome size (Figure 3) and in gene gains along terminal branches between host lifestyles. Data for the other five species, which show similar trends, is presented in Supp. Fig. 4.

We further found that the difference between gene gain and loss counts across all branches is strongly correlated with the number of HGT events per branch, supporting HGT as the primary driver of gene acquisition in these lineages (Fig. 4B). Additionally, reconciliation-based reconstructions of gene copy number along the species tree revealed that increases in gene content predominantly occur along terminal branches associated with industrialized hosts (Fig. 4C and Supp. Fig. 4). Notably, gene families consisting of singletons or occurring in less than 4 StGBs were excluded from this analysis, indicating that proteome expansion is not solely driven by the acquisition of rare genes, but also involves genes that are more broadly shared across strains.

Altogether, these results show that the expansion of bacterial proteomes in industrialized hosts is a relatively recent phenomenon, likely associated with the emergence of industrialized lifestyles, and is mainly driven by high occurrence of HGTs.

### Differential enrichment of genes based on host industrialization status

Building on our finding that host industrialization influences pangenome size and fluidity, we hypothesized that specific gene families within the accessory genome may exhibit differential prevalence between strains from industrialized and non-industrialized hosts. To test this, we analyzed gene presence/absence patterns across the pangenome of each species (mean = 2,691 gene families; SD = 1,515), using phylogeny-aware logistic regression models and the recombination-aware StGB phylogenies. We considered host industrialization status as a binary response variable, and significant associations were further validated using PC1 Lifestyle as a continuous response variable (see Fig. 1B and Methods) for host industrialization. On average, we found that 5.13% (SD = 8.32%) of gene families were significantly associated with host lifestyle (FDR-adjusted p < 0.05) (Fig. 5A, Supp. Fig. 5 & Supp Table 5).

**Figure 5.**
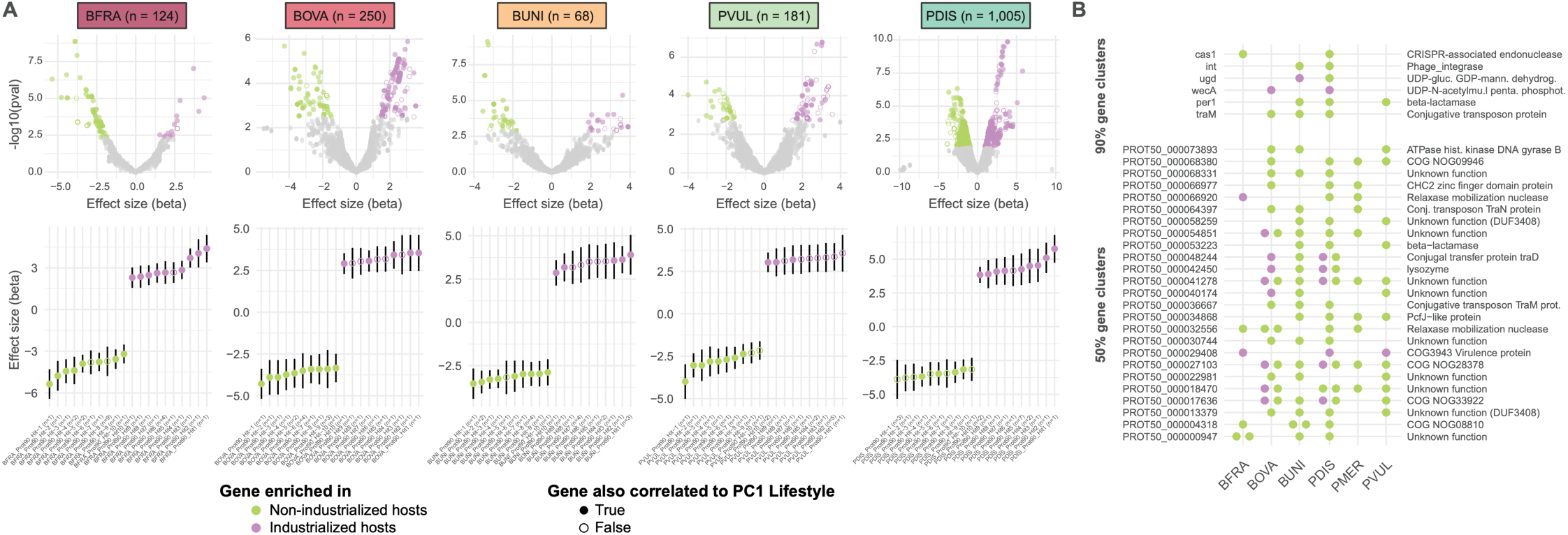
Genes differentially enriched between host industrialized and non-industrialized lifestyles. A. Gene enrichment analysis based on categorical and continuous levels of industrialization. Gene profiles were coded as presence/absence data and were correlated to host industrialization status encoded as a binary variable. Differential enrichment was tested while controlling for phylogeny. Significant hits (q-value < 0.05) are colored coded based on host lifestyle (purple: industrialized; green: non-industrialized). Non-significant genes are colored in grey. Top row: volcano plots showing all genes. The number of statistically significant genes is shown next to each species acronym. Significantly differentially enriched genes validated by measuring correlations with PC1 Lifestyle rather than industrialization status as a binary variable are shown in plain circles. Significant genes not validated with PC1 Lifestyle are shown as empty circles. Bottom row: top 10 most differentially enriched genes, for each lifestyle category. Genes with similar presence/absence profiles are collapsed into a single gene cluster. Gene labels indicate the number (n) of 90% gene families collapsed together. Data for 5 species are shown. These species harbor most of the significant hits. Data for the other 5 species is shown in Supp. Fig. 5. B. Gene families (90% and 50% similarity gene clusters on top and bottom panels, respectively) with signals of differential enrichment across multiple species. Most gene families show convergent signals of differential enrichment based on host lifestyle (enrichment in one of two lifestyle categories across multiple species).

To assess whether gene-level associations reflect convergent adaptation across species, we identified genes with significant industrialization-associated prevalence in multiple taxa. Such convergent signals, replicated both within and between species, would support the hypothesis of positive selection acting on specific genes in response to lifestyle-related environmental pressures. We found six gene families with significant associations in two or more species, with all but one gene showing consistent direction of enrichment direction in relation to industrialization: cas1 (BFRA, PDIS), int (BUNI, PDIS), ugd (BUNI, PDIS), wecA (BOVA, PDIS), per1 (BUNI, PDIS, PVUL), and traM (BOVA, BUNI, PDIS) (Fig. 5B). Interestingly, both ugd and wecA are involved in the formation of bacterial cell surface structures, particularly in the synthesis of polysaccharides and glycans that form key components of the envelope, such as capsule, O-antigen LPS, and enterobacterial common antigens ^45,46^.

Expanding to functional annotations, we identified 23 KEGG Ortholog (KO) groups with significant prevalence associations in at least two species. Of these, four appeared in three or more species: K01185 (lysozyme; BOVA, BUNI, PDIS); K07154 (serine/threonine-protein kinase HipA; BFRA, PDIS, PVUL); K17836 (beta-lactamase class A; BUNI, PDIS, PVUL); and K21572 (starch-binding outer membrane protein SusD/RagB; BFRA, BOVA, BUNI, BXYL, PDIS, PDOR) (Supp. Table 5).

To further explore cross-species convergence, we clustered the species-level catalogs of 90%-similarity gene families into a broader, cross-species catalog at 50% sequence similarity. This revealed 25 gene families with industrialization-associated signals in multiple species (Fig. 5B & Supp. Table 5). Although many of these lacked annotations, we found that four belong to the tra operon, involved in conjugative transfer of genetic material (traD, traK, traM, traN) ^47^, with traK found to be associated with industrialization status in five species (BOVA, BUNI, PDIS, PVUL, PMER). Additionally, two gene families were annotated as relaxases, enzymes involved in the mobilization of plasmids and other mobile elements. Other annotated genes included a putative virulence factor and a lysozyme, both of which may play roles in microbial competition or host interactions (Fig. 5B).

Together, these findings highlight the key role of horizontal gene transfer and mobile genetic elements in shaping the accessory genome in response to host lifestyle ^48,49^. The recurrence of industrialization-associated gene families across multiple species suggests that adaptation to industrialized and non-industrialized environments may be mediated, in part, by the acquisition and spread of genes involved in cell surface structure, conjugation, resistance, and species-species interactions.

### Lifestyle-specific signals of selection at the genome and gene levels

We next quantified gene-level signatures of selection to identify protein-coding genes under positive selection in industrialized or non-industrialized host environments. To do so, we focused on unicopy core protein-coding genes and calculated Ka/Ks ratios separately for strains from each lifestyle category. On average, 1,724 genes were included in the Ka/Ks analysis per species (range: 1,146 for *Bacteroides xylanivorans* to 2,065 for *Bacteroides ovatus*), with 87% of these gene families having sufficient synonymous divergence (Ks) to be analyzed with confidence (see Methods, Supp. Table 6).

Across species, 1.4% of genes showed evidence of positive selection (Ka/Ks > 1) in at least one lifestyle category (Fig. 6A). The prevalence of positively selected genes varied substantially between species, being highest in *P. dorei* (PDOR, 6.05%), *P. merdae* (PMER, 1.89%), and *P. vulgatus* (PVUL, 1.81%), and lowest in *B. ovatus* (BOVA, 0.22%), *B. xylanivorans* (BXYL, 0.31%), and *P. distasonis* (PDIS, 0.36%) (Fig. 6C). These differences are consistent with species-specific variation in overall selection pressure, as reflected in median Ka/Ks values across genes, with PDOR displaying the highest (0.28) and BXYL the lowest (0.09) (Fig. 6B). We also identified numerous genes that were under positive selection specifically in either industrialized or non-industrialized hosts, suggesting context-specific adaptive pressures (Fig. 6A & C-D).

**Figure 6.**
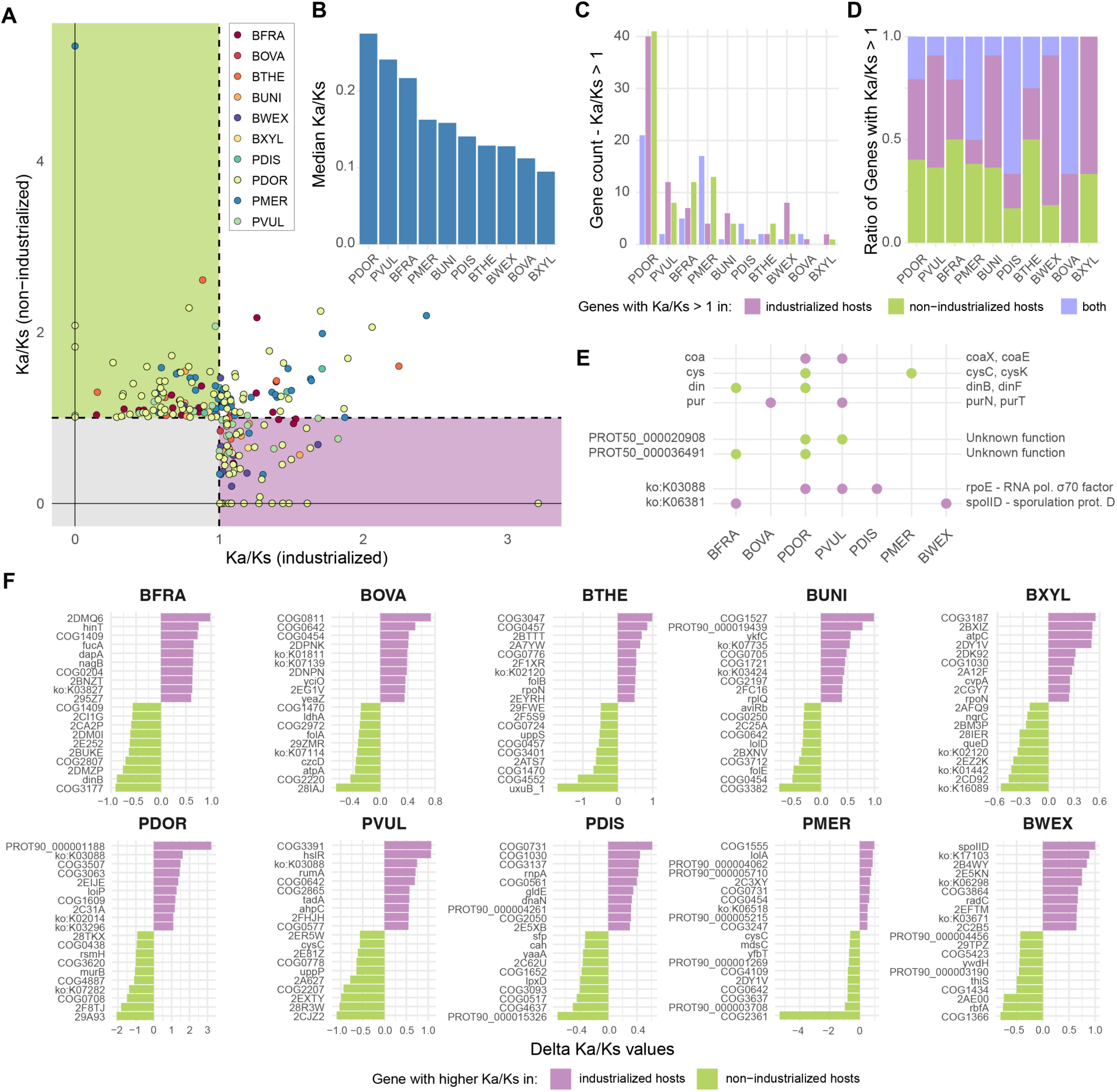
Lifestyle-specific signals of positive selection at the gene level. A. Gene-level Ka/Ks values across species and host lifestyles (90% similarity gene families). For each lifestyle category and each gene, Ka/Ks values were computed for all pairs of codon-aligned gene sequences. Median Ka/Ks values are reported. B. Distribution of species-level median Ka/Ks values, aggregated across all genes. C. Counts of genes with median Ka/Ks values >= 1 (positive selection) across host lifestyle categories. D. Percentage of genes with median Ka/Ks values >= 1 across host lifestyle categories. E. Operons, 50% similarity gene clusters and KEGG KOs with convergent signals of positive selection across 2+ species. F. Top genes with highest absolute differences in median Ka/Ks values between industrialized and non-industrialized lifestyle categories.

To collect additional evidence of positive selection associated with industrialization status, we searched for convergent signals across species – specifically, gene functions under industrialization-specific positive selection in multiple taxa. This analysis revealed two KEGG KOs with repeated signatures of positive selection in strains from industrialized hosts (Fig. 6E). The first, K03088, encodes rpoE, an extracytoplasmic function sigma factor from the sigma70 family, which regulates the expression of stress response genes involved in maintaining cell envelope integrity and responding to oxidative stress ^50^. K03088 genes are under positive selection in strains of PDIS, PDOR, and PVUL in industrialized hosts, suggesting a lifestyle-associated diversification of envelope stress responses in industrialized environments. The second, K06381, groups genes with peptidoglycan lytic transglycosylase activity, including spoIID that is involved in sporulation (Fig. 6E) ^51^. Two genes within this orthologous group showed signals of positive selection in industrialized hosts: spoIID itself in BWEX, and a SpoIID/LytB domain-containing protein in BFRA. These findings suggest that peptidoglycan hydrolase functions – which play roles in cell wall remodeling during sporulation in BWEX – may be adaptive targets in industrialized gut environments.

We also identified six operons containing multiple genes under positive selection across species: genes of the CoA biosynthesis (coa) (coaE in PVUL and coaX in PDOR) and of the purine metabolism (pur) (purT in BOVA and purN in PVUL) operons are under positive selection in industrialized hosts. Genes of the DNA damage response (din) (dinB in BFRA and dinF in PDOR) and of the cysteine metabolism (cys) (cysK in PDOR and cysC in PMER) operons are under positive selection in non-industrialized hosts (Fig. 6E).

To further investigate host lifestyle-driven selection, we ranked genes based on the difference in Ka/Ks between strains from industrialized and non-industrialized hosts, regardless of whether Ka/Ks exceeded 1 (Fig. 6F). This analysis revealed a set of genes with consistently elevated selection pressure in one lifestyle category across multiple species (Supp. Fig. 6), many of which are associated with stress response, metabolism, or virulence (Supp. Fig. 6). For instance, both pepP and tpiA exhibited elevated Ka/Ks in industrialized hosts across eight species. pepP encodes Xaa–Pro aminopeptidase P, a cytosolic metallo-exopeptidase involved in peptide degradation, outer membrane vesicle (OMV) production, and bacterial virulence ^52^. tpiA encodes triosephosphate isomerase, a central enzyme in glycolysis essential for growth on glucose and other glycolytic substrates ^53^. In several bacteria, tpiA has also been linked to virulence, pathogenicity, and antibiotic resistance ^54^. Another gene, truB, showed elevated Ka/Ks in industrialized hosts across seven species. truB encodes a tRNA pseudouridine synthase, which contributes to ribosome stability and stress adaptation. truB has also been implicated in bacterial virulence in pathogenic contexts ^55^.

Conversely, uppS and serC exhibited elevated Ka/Ks in non-industrialized hosts across eight and seven species, respectively (Supp. Fig. 6). uppS encodes an undecaprenyl diphosphate synthase, an essential enzyme in the biosynthesis of lipid carriers required for peptidoglycan synthesis, cell wall assembly, and antibiotic resistance ^56^. serC encodes a phosphoserine aminotransferase, which is critical for serine biosynthesis and supports metabolic flexibility, particularly under nutrient-limited conditions ^57,58^.

Together, these results demonstrate that host industrialization status can exert selective pressures on specific genes and pathways across multiple gut bacterial species.

### Parallel SNV-level adaptation to host industrialization across gut bacteria

Next, we identified single nucleotide variants (SNVs) that are associated with host industrialization status to go beyond the detection of positive selection at the level of genes, as individual variants may also evolve in response to host environments. To investigate this, we analyzed non-synonymous SNVs in the core genome, while accounting for phylogenetic structure (see Methods). To limit the confounding effect of recombination, we excluded SNVs predicted to be part of recombination regions ^59^ (see Methods). This analysis was restricted to the five species with the largest number of genomes and StGBs (BFRA, BOVA, BUNI, PDIS and PVUL) to ensure sufficient statistical power for detecting lifestyle-associated SNVs. To validate associations drawn from the analysis of isolate genomes, we performed a secondary association analysis using SNVs called from MAGs reconstructed from gut shotgun metagenomes of the entire GMbC cohort (n = 1,015 individuals, 12 countries and 35 localities spanning a range of a range of industrialization levels and subsistence strategies), which we recently described ^6^ (see Methods). We also considered an additional layer of validation, using PC1 Lifestyle as a continuous response variable for host industrialization. We then defined an SNV hit as “high-confidence” if (i) it showed a consistent direction of effect with both isolate and MAG data, (ii) it achieved FDR-adjusted significance (p-val < 0.05) in the isolate genome analysis, (iii) it achieved significance (p-val < 0.05) in the validation analysis with MAGs, and (iv) it achieved significance (p-val < 0.05) in the validation analysis with PC1 Lifestyle (Supp. Table 7).

Across the five species tested, we found numerous high-confidence non-synonymous SNVs in core genes that are significantly associated with host industrialization (Fig. 7A). The identification of such SNVs suggests that convergent adaptation to host industrialization across distantly-related strains is occurring within each of these species. Interestingly, 93%, 71%, 79%, 85% and 100% of hits detected with industrialization categories were validated with PC1 Lifestyle for BFRA, BOVA, BUNI, PDIS and PVUL, respectively. Beyond showing robustness, these validations also suggest that changes in non-synonymous SNV frequencies occur along a spectrum of host lifestyle variation. Interestingly, genes containing these host lifestyle-associated SNVs are part of operons involved in stress response (e.g. mut, uvr), nutrient acquisition and central metabolism (e.g. asn, nad), host interaction and virulence (e.g. ent, prt), and transport and membrane remodeling (e.g. feo, mal) (Fig. 7A & Supp Table 7).

**Figure 7.**
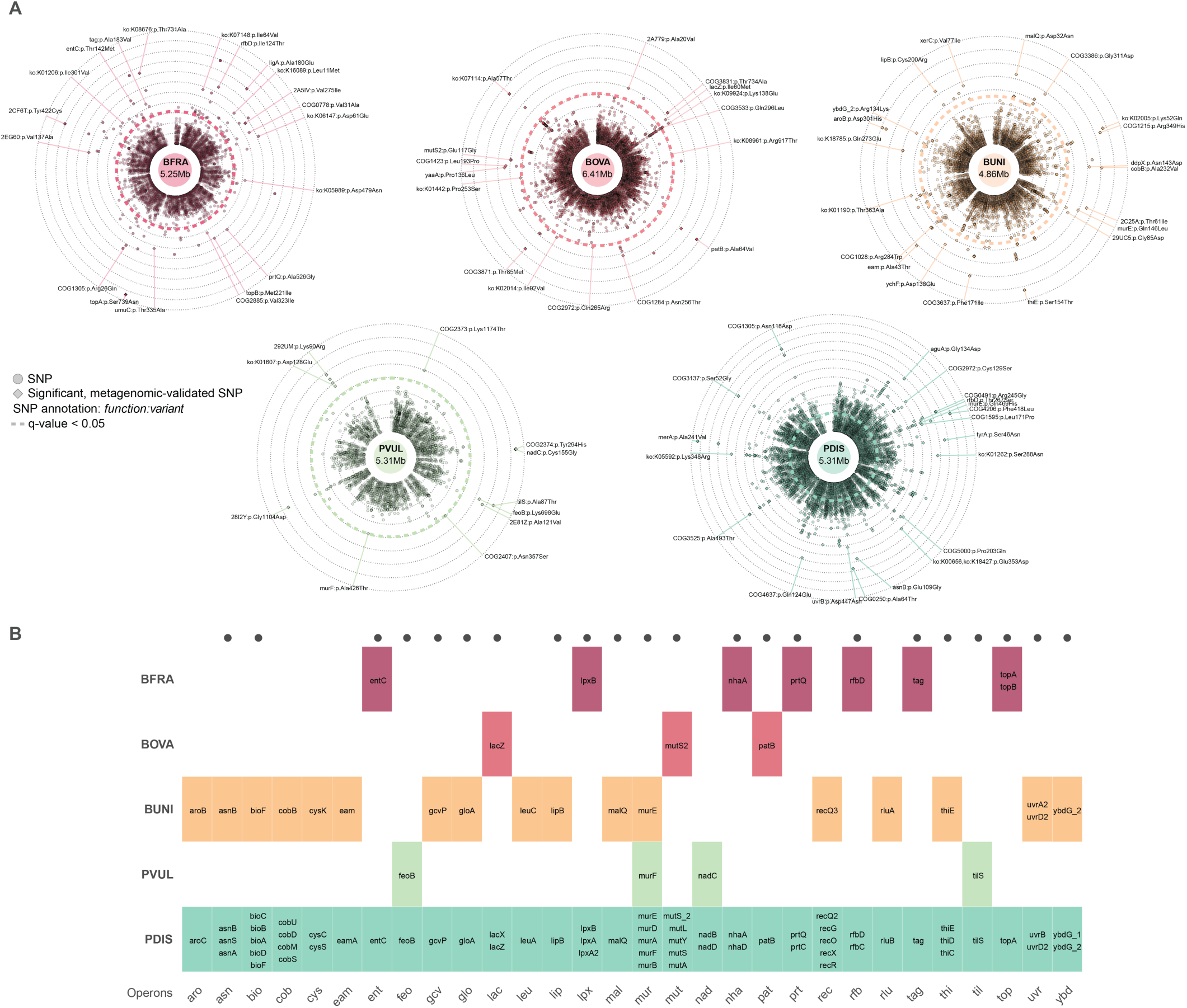
Patterns of single nucleotide polymorphisms reveal genes and operons that undergo cross-species parallel evolution associated with host lifestyle. A. Associations between single nucleotide variants (SNVs) and host lifestyle categories were calculated while accounting for phylogeny (see Methods). Associations were calculated for the 5 species with enough genome sample size to yield sufficient statistical power. Hits (q-value < 0.05) were cross-validated using GMbC shotgun metagenomes (n = 1,015, see Methods) that were sampled from diverse geographies worldwide, including those from which isolate genomes originate (reference). Hits validated by metagenomes are shown as diamonds and annotated. B. Convergent signals of SNV-host lifestyle associations across bacterial species. Each tile represents an operon–species association and contains the names of genes within that operon that contain host-lifestyle associated SNVs. Operons are shown along the x-axis, and species along the y-axis, cells are color-coded by species identity. Black points show operons in which similar genes contain host-lifestyle associated SNVs across species. All SNV data can be found in Supp. Table 7.

We then searched for genes and gene functions that harbor lifestyle-associated non-synonymous SNVs in multiple species. As for our previous analyses on gene enrichment and Ka/Ks, the rationale is that cross-species convergent evolution would provide even further compelling evidence of adaptation to host environments shaped by industrialization beyond strain-level convergent evolution. This analysis revealed 23 genes with lifestyle-associated non-synonymous SNVs occurring in at least two different species (Fig. 7B). We further identified 30 operons containing genes with lifestyle-associated non-synonymous SNVs across multiple species. Remarkably, we found that the cob operon, which encodes enzymes for cobalamin (vitamin B12) biosynthesis, harbors lifestyle-associated SNVs in PDIS and PUNI. This mirrors recent findings from our analysis of GMbC gut metagenomes ^6^, which identified cobalamin biosynthesis pathways as significantly enriched in populations having increased levels of industrialization, after adjusting for geographic, dietary, and host genetic confounders. These two independent lines of evidence, found at the genomic and metagenomic levels, strongly suggest that vitamin B12 biosynthesis is a key adaptive trait in response to industrialized host environments. Second, we observed repeated signals in the mur operon, which is essential for murein (peptidoglycan) biosynthesis, a core component of the bacterial cell wall. Specifically, lifestyle-associated non-synonymous SNVs were identified in murE in BUNI, murF in PVUL, and murA, murB, murD, murE, and murF in PDIS. This points to a potential role of cell wall remodeling and integrity in adaptation to differing host lifestyles. Third, we found that two operons involved in lipopolysaccharide (LPS) biosynthesis (lpx, encoding genes for lipid A biosynthesis and rfb, encoding genes for O-antigen biosynthesis) also harbor host lifestyle-associated SNVs in multiple species. Given the role of LPS in host immune recognition and membrane structure, these findings suggest that modulation of LPS structure may be an important contributor to adaptation to host lifestyle-specific environments.

Together, these findings provide evidence for SNV-level parallel evolution across gut bacteria, highlighting shared functional targets that may mediate adaptation to host lifestyle and industrialization.

## Discussion

In this study, we provide evidence that human lifestyle, particularly factors associated with industrialization status, shape the genomic evolution of bacterial commensals. We show that strains from industrialized hosts harbor larger proteomes and exhibit increased pangenome fluidity. We found that recently acquired genes during bacterial evolution, explained by HGT, are driving these phenomena. These findings confirm that HGT is a primary mechanism for rapid evolutionary innovation in the human gut microbiome, and are in line with our previous discovery that frequencies of HGT are elevated among microbes co-occurring in industrialized hosts ^21^. Interestingly, a recent study also showed that HGT facilitates adaptative selective sweeps of *mdxEF* genes, involved in maltodextrin metabolism, among industrialized populations ^48^. We further accumulated multiple lines of evidence of convergent evolution of genomic features with respect to host industrialization status and lifestyle (PC1 Lifestyle), including the differential enrichment of genes, gene-level positive selection, and non-synonymous SNV frequency. Such convergent patterns could be observed at different taxonomic resolutions, both between strains and across species. These parallel genomic signatures are consistent with adaptation to the ecological conditions associated with industrialized or non-industrialized environments, including exposure to different commensal and environmental microbes, dietary profiles and medication usage ^60^. The functional categories involved in these changes include traits that are relevant for ecological adaptation. In particular, genes under selection or containing lifestyle-associated SNVs are involved in stress response (e.g., rpoE, uvr, mut), cell envelope remodeling (e.g., mur, lpx, rfb), central metabolism (e.g., cob, nad, asn), and host interaction or virulence-related functions (e.g., prt, ent) (Fig. 5 & 6).

Lastly, several of the adaptive features that we identified may have important implications for host physiology and health. Notably, genes involved in vitamin B12 biosynthesis harbor lifestyle-associated non-synonymous SNVs across multiple bacterial species. Several vitamin B12 biosynthesis pathways were also found to be differentially abundant in the metagenome of the broader GMbC cohort (n = 1,015) ^6^, with higher abundance among populations more exposed to industrialized environments. This suggests that cobalamin biosynthesis constitutes a key adaptive trait in industrialized gut ecosystems. Similarly, we observed multiple signals of adaptation in lipopolysaccharide (LPS) biosynthesis pathways – including wecA, lpx, and rfb genes (Figs. 5 & 7) – which are critical for bacterial membrane structure, bacteria-bacteria interactions and host immune recognition. These findings indicate that LPS remodeling may not only contribute to microbial adaptation in industrialized hosts, but could also influence host immune and inflammatory responses ^61^.

### Limitations

Our analysis focused on ten bacterial species. While these species are prevalent and functionally significant components of the human gut microbiome, further investigations that broadens taxonomic and host representation will be necessary to assess the generalizability of our findings. In addition, even though we discovered several associations between host industrialization status and microbial genomic features, it is still unclear which environmental factors, such as dietary composition, medication use, sanitation, or other exposures related to lifestyle, are responsible for these correlations. Future research combining functional validation, experimental evolution, and fine-scale environmental variable measurement will be essential to identify and separate the selective pressures that influence the evolution of microbial genomes in response to variations in the industrialization of host environments.

Finally, the evolutionary mechanisms driving the expansion of proteomes in strains from industrialized hosts remain unclear. Although we find signatures of adaptation shaping the distribution of individual genes, the overall proteome expansion may result from the relaxation of purifying selection, allowing the accumulation of slightly deleterious genes and mobile elements. Future studies involving population structure reconstruction, effective population size estimation, the study of mutational and recombination processes influencing allele frequencies, and fitness experiments will be required to distinguish the relative contributions of adaptive versus non-adaptive processes in influencing the evolution of the bacterial genome across lifestyles.

## Methods

### Participant recruitment and collection of biospecimens

To culture and isolate the additional 1,841 GMbC bacterial strains generated in this study, we selected 20 participants from the GMbC cohort (cross-sectional population cohort of healthy adults that we recently described ^6^). These individuals were chosen to represent a broad range of lifestyle, urbanization levels, and geographic origins (see below), and none had taken antibiotics or antiparasitic treatments recently. All participants were asymptomatic for infectious or chronic diseases at the time of enrollment. The 20 participants were from Cameroon, Ghana, Malaysia, Nigeria, and Rwanda. This new set of isolate genomes builds upon our initial GMbC collection of 4,140 isolates ^21^, bringing the total to 6,000 isolates from 56 participants spanning nine countries, including the USA, Canada, Finland, and Tanzania. Written informed consent was obtained from all participants, using translations in local language when appropriate. Research & ethics approvals were obtained from the MIT IRB (protocol #1612797956) and from the Ethics commission of the Medical Faculty of Kiel University (Studie D 511/24). Permits were also obtained in each sampled country prior to the start of sample collection, from the following ethics committees:

- Cameroon: Comité National d’Ethique de la Recherche pour la Santé Humaine, protocol #2017/05/901/CE/CNERSH/SP;
- Ghana: Cape Coast Teaching Hospital Ethical Review Committee, protocol #CCTHERC/RS/EC/2016/3 and Committee on Human Research, Publication and Ethics of the Komfo Anokye Teaching Hospital, protocol #CHRPE/AP/398/18;
- Malaysia: Universiti Malaya Medical Research Ethics Committee, MREC ID No.: 2018219-6033;
- Nigeria: National Health Research Ethics Committee of Nigeria, protocol #NHREC/01/01/2007-29/04/2018.
- Rwanda: National Ethics Committee, protocol IRB 00001497 of IORG0001100

Participants self-collected a fresh fecal sample using sterile containers, which were returned to GMbC scientists for on-site processing within 3 hours after defecation. Raw stool was diluted 1:5 in a 25% pre-reduced anaerobic glycerol solution containing acid-washed glass beads for homogenization, then aliquoted into 2 mL cryogenic tubes. Aliquots in cryoprotectant were flash-frozen either in liquid nitrogen (−196 °C) using a cryoshipper tank or stored at −80 °C. An additional 1–2 g of stool was preserved in RNAlater for DNA stabilization and sequencing. All samples were subsequently shipped to MIT for further processing and storage.

### Host metadata and industrialization status

Lifestyle metadata were collected for all GMbC participants and analyzed using dimensionality reduction techniques ^6^. Briefly, we used the Human Development Index (HDI), as described previously ^21^, as a measure to determine the industrialization status of sampled populations. Populations from localities with HDI values below the national median (0.739 in 2022) were classified as non-industrialized, while those with values above this threshold were classified as industrialized. We also collected a range of lifestyle variables including, e.g., population density, main subsistence strategies, type of floor in households, access to electricity, household size, contact with animals, frequency of physical activities per week and source of drinking water. A full list of lifestyle variables, along with dietary and medication data for the participants used for bacterial culturing is provided in Supp. Table 1.

We used the PCAmix function from the PCAmixdata R package to perform a principal component analysis of lifestyle variables (Fig. 1B). PC1 Lifestyle strongly correlates with industrialization status, along with other key factors such as access to electricity, main subsistence strategy, floor type and type of drinking water (Fig. 1B). PC1 Lifestyle is used as a continuous proxy for industrialization.

### Culturing and isolation of gut bacteria

We used the same culturing procedures as used previously to build the BIO-ML ^12^ and the GMbC ^21^ isolate collections. Fecal samples were processed anaerobically (5% Hydrogen, 20% Carbon dioxide, balanced with Nitrogen) and diluted in pre-reduced PBS (with 0.1% L-cysteine hydrochloride hydrate). Samples were plated onto pre-reduced agar plates and incubated anaerobically at 37C for 7 to 14 days. Both general (nonselective) and selective media were used to culture diverse groups of organisms (Supp Table X). After incubation, bacteria were isolated and purified through two rounds of picking and re-streaking at 37C. One colony was then inoculated in liquid media. After 2 days of anaerobic incubation at 37C, the taxonomy of the isolate was identified using 16S rRNA gene Sanger sequencing (starting at the V4 region). We first amplified the full 16S rRNA gene by PCR (27f 50-AGAGTTTGATCMTGGCTCAG-30 - 1492r 50-GGTTACCTTGTTACGACTT-30) and then generated a 1kb long sequence by Sanger reaction (u515 50-GTGCCAGCMGCCGCGGTAA-30). Isolates were then stored at -80C in a pre-reduced cryoprotectant glycerol buffer. See Supp Table X for the list of culturing media, selection treatments and isolate metadata.

### DNA extraction, library prep and sequencing of isolate genomes

We used the same procedures used to generate the first set of GMbC isolate genomes ^21^. Briefly, whole genome DNA from isolates was extracted with the DNeasy UltraClean96 MicrobioalKit (QIAGEN), following manufacturers’ protocols. Genomic DNA libraries were built from 1.2ng of extracted DNA using the Nextera DNA Library Preparation kit (Illumina), following the manufacturer’s protocol, with reaction volumes scaled accordingly. Isolated libraries were pooled, with insert size and concentration of each pooled library being determined using an Agilent Bioanalyzer DNA 1000 kit (Agilent Technologies). Paired-end (2×150bp) reads sequencing was performed using an Illumina NextSeq 500 instrument (Illumina Inc) at the Broad Institute.

### Assembly and quality estimation of GMbC isolate genomes and MAGs

Isolate genome assemblies were reconstructed as follows. We first used cutadapt v1.12 ^62^ (with parameters -a CTGTCTCTTAT -A CTGTCTCTTAT) and Trimmomatic v0.36 ^63^ (with parameters PE -phred33 LEADING:3 TRAILING:3 SLIDINGWINDOW:5:20 MINLEN:50) on demultiplexed paired-end short reads to remove barcodes and Illumina adapters and for base quality filtering. We then used SPAdes v.3.9.1 ^64^ (with parameter–careful) for de novo assembly of reads into contigs. To iteratively improve genome assemblies, we used SSPACE v3.0 ^65^ and GapFiller v1-10 ^66^ to scaffold contigs and to fill sequence gaps (with default parameters). We removed scaffolds smaller than 1kb. All reads were aligned back to the assembly to compute genome coverage using BBmap v37.68 (https://jgi.doe.gov/data-and-tools/bbtools/) and the covstats option (with default parameters).

MAGs were reconstructed from n = 1,015 shotgun metagenomes of the GMbC cohort, as described in our recent work ^6^, resulting in a total of n = 24,163 quality-filtered MAGs (see below). Among these, 1,424 MAGs originated from the donors used for generating GMbC isolate genomes (from the same original stool sample). Briefly, quality filtered short read shotgun metagenomic data was assembled into contigs using Megahit ^67^. Contigs were binned using a multi-binning and refinement strategy in MAGScoT ^40^, using MaxBin2 ^68^, Metabat2 ^69^, CONCOCT ^70^ and VAMB ^71^ binning tools.

Genome completeness and contamination were estimated using CheckM and MAGScoT ^40,72^. Standard thresholds for completeness (>50%) and contamination (<10%) were used to filter out low-quality genomes. Genome quality was defined as Q = Completeness - 0.5 * Contamination. We considered genomes with Q > 0.7 as high-quality genomes, and genomes with 0.7 > Q > 0.5 as medium-quality genomes. All GMbC isolate genomes are high-quality genomes, with median Q = 0.98, min Q = 0.88, max Q = 1. Assembly and genome quality statistics for GMbC isolate genomes and MAGs can be found in Supp. Table 1.

### External collections of human gut bacterial isolate genomes and MAGs

We included 3,632 human gut bacterial isolate genomes from the BIO-ML collection, which we generated in a previous study ^12^. These isolates were obtained from participants with industrialized lifestyles recruited in the Boston (MA, USA) area and were processed using the same culturing, sequencing, and assembly protocols described above for the GMbC isolate genomes. MAGs were also reconstructed from the 11 BIO-ML participants used for culturing and isolating bacteria, using the same pipeline as for GMbC MAGs. For comparative analyses, we also included 4,644 representative MAGs from the UHGGv2 collection of human gut MAGs ^24,36^.

### GMbC shotgun metagenomes

We generated shotgun gut metagenomic data for GMbC cross-sectional population cohort (n = 1,015 healthy adult participants). This includes the participants whose sample we used in this study to culture and isolate gut bacteria. We present the shotgun metagenomic data in our recently published work ^6^. Briefly, the data were sampled from 12 countries and 35 localities, across a range of industrialization levels and subsistence strategies. Briefly, DNA was extracted from stool samples stored in RNAlater using the MoBio Powersoil 96 kit, and sequencing libraries were prepared with the Nextera XT kit. Paired-end 150 bp sequencing was performed on the Illumina NovaSeq S4 platform, generating a median of 21.7 million reads per sample. GMbC and BIO-ML MAGs were reconstructed as described above (section “Assembly and quality estimation of GMbC isolate genomes and MAGs”). GMbC and BIO-ML MAGs and isolate genomes were all clustered into reference species-level genome bins (SGBs) using a multi-step workflow based on ANI clustering with dRep and fastANI, resulting in a final non-redundant set of 2,379 SGBs, 480 of which contained isolate genomes ^6^. SGB representatives were taxonomically annotated with GTDB-Tk (v2.3) and the GTDB reference database (rel214). They were used as references for abundance estimation with Salmon in metagenome mode ^73^. Relative abundances were expressed in TPMs, filtered to remove low-abundance SGBs (<1,000 reads and <250 TPMs), and transformed using CLR with a pseudocount of 1. Fig. 1B shows the microbiome alpha and beta diversity of GMbC and BIO-ML participants from whom bacterial isolates were obtained. Alpha diversity was calculated using Faith’s PD index, while beta diversity was assessed using the unweighted UniFrac dissimilarity implemented in the GUniFrac R package (https://cran.r-project.org/web/packages/GUniFrac/index.html). Both metrics were computed using a phylogenetic tree of the reference SGBs based on the GTDB single-copy marker gene alignments and the GTDBtk “infer” workflow.

### SGB and StGB reconstruction from isolate genomes and MAGs

We clustered genomes into species-level genome bins (SGBs) by combining GMbC and BIO-ML isolate genomes with GMbC MAGs. To ensure high-quality clustering, we implemented an iterative workflow that optimizes cluster resolution and quality. First, GMbC MAGs were clustered within each geographic location at 97% average nucleotide identity (ANI) using dRep (v3.4.0). For each cluster, we selected the MAG with the highest quality score as its representative. These representative MAGs were then classified as either high-quality (HQ; score > 0.7) or medium-quality (MQ; 0.5 ≤ score ≤ 0.7). Next, HQ MAG representatives were combined with all GMbC and BIO-ML isolate genomes and clustered into SGBs (95% ANI). Each MQ MAG representative was then compared to the HQ SGB representatives using fastANI (v1.33) and assigned to an SGB if it shared at least 95% ANI. MQ MAGs without matches at the required threshold were reclustered using dRep to form MQ SGBs, and we conserved bins only if they contained more than one genome. Finally, HQ and MQ clusters were merged to generate the final set of non-redundant SGBs. GMbC and BIO-ML isolate genomes were also clustered at 99% ANI to form strain-level genome bins (StGBs). For the evolutionary genomic analyses of the 10 species presented in Fig. 3-7, StGBs containing genomes from multiple donors were further split by donor to ensure that each StGB represented a donor-specific strain.

### Comparison of GMbC isolates with UHGG MAGs, BIO-ML isolates and GMbC MAGs

SGB and StGB representative genomes were compared to the UHGGv2 representative MAGs (n = 4,644) using fastANI. A genome was classified as “included in UHGG” if it shared >95% ANI with a UHGG representative genome. GMbC isolate genomes were considered to be represented in GMbC MAGs or BIO-ML isolates if they clustered within the same SGB or StGB.

### Identification of Isolate-MAG pairs of genomes

Isolate-MAG pairs from the same donor and same species-level taxonomy were identified by matching MAGs and isolate genomes from the same individual that clustered within the same SGB. If multiple isolate genomes from a given donor were present within an SGB, we calculated pairwise ANI values between each isolate and the corresponding MAG. The isolate genome with the highest ANI to the MAG was selected as the representative genome for downstream pairwise comparisons of genomic features (Fig. 2).

### Gap-seq metabolic features

Genome-scale metabolic models were reconstructed using gapseq (v1.2, commit 2dfa8c80) ^74^ for all paired MAGs and isolate genomes (Fig. 2). The reconstruction workflow using gapseq consisted of 5 steps. First, all bacterial pathways from the MetaCyc and gapseq database were retrieved with ‘gapseq find’ and options ‘-p all -t Bacteria’. Second, all cross-membrane metabolite transporters were recovered using ‘gapseq find-transport’. Third, draft metabolic networks were reconstructed from the two previous steps using ‘gapseq draft’. Fourth, an anoxic growth medium for the given organism is predicted from the draft metabolic network using ‘gapseq medium’ and the option ‘-c "cpd00007:0"’, following our previous study ^75^. Finally, gaps in the draft metabolic network were filled in order to enable growth (i.e. formation of biomass), assuming the growth medium predicted at the previous step. This step was performed using the ‘gapseq fill’ module. For all models, the number of reactions, metabolites, and genes that are linked to reactions or transporters are counted for the draft network reconstruction and for the gap-filled network.

### Functional profiling of ARGs, VFs, CAZymes, MGEs and MGE machineries in isolate-MAG pairs

Antibiotic resistance genes and virulence factors were identified in genome assemblies using Abricate (https://github.com/tseemann/abricate), with annotations based on the NCBI AMRFinderPlus database ^76^ for resistance genes and the VFDB database ^77^ for virulence factors. CAZyme genes were detected using dbCAN3 (https://github.com/linnabrown/run_dbcan) ^78^. We used MacSyFinder (version 2.0) ^79^ to profile MGEs and MGE machineries of three key systems: type secretion systems (TXSS), type IV filaments (TFF), and conjugative transfer systems (Conj). We used ISEScan (version 1.7.2.3) ^80^ to identify insertion sequences (IS). We used geNomad (v1.7.4) ^81^ to detect prophages. Following a recently established protocol ^82^, we first used checkV (v1.5) ^83^ to measure phage sequence quality, retaining only prophage sequences with >50% completeness (including medium-quality, high-quality and complete sequences) for downstream analysis. Prophage sequences were then clustered based on average nucleotide identity, with the longest prophage contig in each cluster selected as the representative sequence.

### Inference of horizontal gene transfers from isolate genomes and MAGs

We detected HGTs on the 147 pairs of MAG and isolate genomes, identifying HGTs among MAGs and isolate genomes separately. We focused on HGTs occurring between bacterial species. The 147 genomes cluster into 87 different SGBs, which constitutes 3,741 species pairs. To detect HGTs, we used methods that we implemented in previous studies ^21,41^ that rely on the Blast-based detection of blocks of DNA that are shared by two genomes of different species. We retained blast hits with 100% similarity and that are larger than 500bp, to focus on the most recent HGTs ^21^. We analyzed HGT sequences that have a relative read coverage higher than 0.2 compared to the average genome coverage in at least one of the two compared genomes ^21^. We considered a HGT having occurred between two species when we could identify at least one genome pair with at least one blast hit.

### Species selection

We ranked species in the combined GMbC and BIO-ML isolate genome collection based on the total number of distinct StGBs sampled, as well as their representation across industrialized and non-industrialized hosts. For downstream genomic analyses, we selected the top 10 species meeting these criteria (Fig. 1G), each represented by a minimum of eight StGBs per host industrialization category (Supp. Table X). Included species are *Bacteroides fragilis* (BFRA), *Bacteroides ovatus* (BOVA), *Bacteroides thetaiotaomicron* (BTHE), *Bacteroides uniformis* (BUNI), *Bacteroides xylanisolvens* (BXYL), *Phocaeicola dorei* (PDOR), *Phocaeicola vulgatus* (PVUL), *Parabacteroides distasonis* (PDIS), *Parabacteroides merdae* (PMER), and *Blautia A wexlerae* (BWEX). We purposefully excluded *Escherichia coli*, as our focus was on abundant members of the gut microbiome that have not yet been extensively characterized – unlike *E. coli*, whose global genomic diversity has been well studied ^84^.

### Variant calling and annotation

For the ten species, we considered all GMbC and BIO-ML isolate genomes with >95% completeness, based on MAGScoT estimates. A reference genome per species was chosen based on the best quality score Q (see above) and the highest L50 value. Contigs were joined into a single fasta sequence with a stretch of 100 ambiguous bases between contigs to create an artificial continuous genome for downstream read mapping and variant calling. Short read data of all isolate genomes were mapped against their respective species reference genome using snippy with default parameters, requiring a coverage >= 10-fold for the variant calling. MAGs derived from metagenomic data of the broader GMbC cohort ^6^ that are of the same species were also included. We selected MAGs that have >97% average ANI (using skani) to any of the isolate genomes in each species, and that have low contamination estimates (<10%) data. MAGs were processed with snippy in the “contigs” mode (--ctgs flag) (https://github.com/tseemann/snippy). Isolate and MAG-derived snippy variant calls were combined in a single whole-genome alignment using the ‘snippy-core’ command. The alignment was then cleaned using the ‘snippy-clean_full_aln’ utility script provided by the snippy developer. To discriminate gaps within the alignment from gaps at the end of contigs, gaps of length <= 10bp were masked. We discarded genomes that had more than 40% ambiguity calls or gaps from the final snippy output. The alignment was converted to a VCF file of all single-nucleotide variants (SNVs) using ‘snp-sites’ utility script of snippy. Multi-allelic variants were decomposed to individual entries in the VCF file using vt decompose (https://github.com/atks/vt). Variant effects were predicted using snpEff ^85^.

### Recombination-aware reconstruction of StGB phylogenies

A reference genome per StGB and per individual was selected using the same criteria as for the selection of species-level reference genomes (see previous section). Reference genomes were extracted from the whole genome alignment reconstructed with snippy and used for phylogeny reconstruction using IQ-TREE2 and a generally time reversible (GTR) substitution model ^86^. We then used Gubbins ^59^ to identify and mask regions of homologous recombination. The masked alignment was then reused in IQ-TREE2 (GTR model) for phylogenetic reconstruction.

### Gene calling and pangenome reconstruction

Protein-coding gene sequences from all isolate genomes of the ten selected species were predicted using PROKKA (default parameters) ^87^. Coding sequences (CDSs) were then clustered into gene families at 90% sequence identity using MMseqs2 (v13.45111) and the ‘easy-linclust’ module with the options ‘--cov-mode 1 -c 0.8 --kmer-per-seq 80’ ^88^. Representative sequences for each gene family were functionally annotated using EggNOG-mapper (v2.1.3; database version 220425) ^89^. Within each species, gene families were subsequently categorized into three groups based on their prevalence across genomes: (soft-)core genome (>95% of genomes), accessory genome (5–95%), and cloud genome (<5%).

### Quantifying mutational selective pressures

Species-level single-copy core gene families (SCGs) were defined as being present in 95% of all isolate genomes of a species, and found in each individual StGB. SCG protein sequences of the representative StGB genomes were aligned using MAFFT in ‘--auto’ mode ^90^. Protein-level alignments were translated back to nucleotidic sequences to obtain codon-level alignments, which were subjected to pairwise Ka and Ks value calculations using the kaks() function of the ‘seqinr’ R package. Sequence pairs with at least one synonymous and one non-synonymous mutation were included, while others were assigned a -1 value and excluded from the analysis as recommended in PAML. For each gene family, mean Ka/Ks values were calculated for each host industrialization category. Mean Ka/Ks values for which > 90% of pairwise comparisons were excluded due to lack of mutations were marked as ‘low confidence’.

### Calculation of the pangenome fluidity

We calculated pangenome fluidity as described previously ^91,92^. Briefly, pangenome fluidity measures the average proportion of genes that are not shared between pairs of genomes from the same species. For each species, we computed fluidity separately for strains isolated from industrialized and non-industrialized hosts, using the representative genome of each StGB. Specifically, we calculated the pairwise proportion of non-shared gene families between all StGB pairs within each lifestyle category. To compare the distribution of non-shared gene proportions between lifestyles, we performed Wilcoxon rank-sum tests.

### Gene tree – species tree phylogenetic reconciliations

We used the species tree-aware gene tree reconciliation method, ALE ^93^ implemented in AleRax ^44^ (commit version: 8705a60, 2025-06-02) to harvest signals of gene evolutionary events (gene duplications, transfers and losses) from all the gene families of the 10 selected species. We used the recombination-aware strain-level StGB phylogenies (see above) as species trees, and all individual gene trees of the gene families containing at least 4 representative genomes. The species tree was midpoint-rooted using ete3 ^94^. The nucleic acid sequences were ordered based on the 90% sequence identity clustering into per-family sequence files and deduplicated (keeping only the gene families which still had at least 4 unique sequences for downstream analysis). The number of gene families per species distributes as follows: BFRA: 4095; BOVA: 6238; BTHE: 4436; BUNI: 4906; BWEX: 3596; BXYL: 5047; PDIS: 5042; PDOR: 2968; PMER: 3340; PVUL: 4988. Gene families were aligned using MAFFT (with L-INS-i option) ^90^ with default settings. From the alignments, gene trees were inferred under the GTR+G+F model using IQ-TREE2 ^86^ and generating 1000 ultrafast bootstrap samples ^95^ with the ‘-bnni’ option. AleRax analysis was performed on these ultrafast bootstrap gene tree samples of all the gene families using the GLOBAL model parametrization (i.e. the DTL-rates are shared among families and jointly estimated), the UndatedDTL reconciliation model and taking 1000 reconciled gene trees along the species tree. Based on the reconciled gene trees, AleRax reports the number of evolutionary events along the branches and leaves of the species tree. Wagner-parsimony ^96^ was used on the species tree to deduce host industrialization status at internal nodes based on existing status at terminal nodes. Evolutionary events could then be associated with industrialization state on a per branch level (Fig. 4A). All reconciliation data can be found in Supp Table 4.

### Gene Family enrichment analysis

Accessory gene families (present in 5–95% of genomes) were included in the gene enrichment analysis if they were found in at least five StGBs and absent from at least five others. Gene distribution data were encoded as presence/absence data. To detect differentially enriched genes while accounting for phylogenetic structure, we used a phylogeny-aware logistic regression model implemented in the phylolm R package ^97^. Host industrialization status (industrialized vs. non-industrialized) was used as a binary explanatory variable, and we additionally tested associations using PC1 Lifestyle, a continuous proxy of industrialization derived from multidimensional lifestyle data (see section above). Significant associations identified with the binary industrialization variable were validated using PC1 Lifestyle and a phylogeny-aware linear regression model in phylolm. Recombination-aware phylogenies at the StGB level were used in all phylolm models. All genomes within each species were included in the regression analyses, with an arbitrary branch length equal to 1% of the median cophenetic distance added to non-representative isolate genomes to preserve tree topology.

### Detection of bacterial single nucleotide variants (SNVs) associated with host industrialization status

For SNV–host industrialization associations, we analyzed biallelic and non-synonymous variants located within single-copy core gene families (defined as present in >99% of genomes), retaining only variants with a minor allele frequency >5%. Variants located in predicted recombination regions were masked prior to analysis. Associations were tested using a logistic regression model that accounts for phylogenetic structure (‘phylolm’ R package). Host industrialization was considered both as a binary response variable (industrialized vs. non-industrialized) and using PC1 Lifestyle as a continuous proxy, consistent with the approach used in the gene enrichment analyses. The latter served as a validation for significant associations identified using the binary variable.

To further validate SNV-level associations, we examined corresponding significant SNVs in the metagenomic data of the broader GMbC cohort by mapping metagenomes against the reference genome of each species (see section “Variant calling and annotation” above). Associations with metagenome-derived SNVs were also tested using logistic regression against host industrialization status as a binary variable.

## Data availability

Short read data and assemblies of GMbC isolate genomes generated in this study will be made available online on the dbGaP server (Study ID: 38715; Accession: phs002235.v1.p1; Accession: phs002205.v1.p1) upon publication of the article. GMbC metagenomes used in this study and published in our recent study ^6^ will be made available at the same dbGaP study.

## Code availability

Scripts and command lines used to process the data will be made available at https://github.com/MMmicrobiome-Lab upon publication of the article.

## List of Supplementary materials

### Supplementary Figures

**Supplementary Figure 1 – Distribution and functional predictions of GMbC genomes reveal host and strain-level diversity.**

A. Distribution of SGB and genome counts across localities and individual hosts. Colors denote country.
B. Phylogenomic tree of representative genomes from 434 species-level genome bins (SGBs). Outer rings show annotations of functional and phenotypic categories predicted by Traitar. For each trait and SGB, trait prevalence was first calculated across all genomes of the corresponding strain-level genome bins (StGBs), and SGB-level conservation scores were then obtained by averaging StGB prevalence values. Trait conservation scores are visualized using an opacity gradient (‘score’ variable).
C. Heatmap of strain-level amino acid auxotrophy variability across species (y axis). Values show the variance in auxotrophy predictions across genomes of a given species. Only genomes with quality score higher than 95% were included in the variance calculations. Auxotrophy predictions were performed from gapseq-derived genome-scale metabolic models (see Methods).

**Supplementary Figure 2 – Pairwise comparisons of genomic features between MAG and isolate genomes sampled from the same donor highlight the recovery limitations of MAGs.**

A. Metagenomic abundance of species included in the MAG vs. isolate genome comparison. Each data point represents a pair of MAG and isolate genome from the same species, sampled within the same host. Species are colored by phylum.
B. Counts of various genomic features in the paired MAG and isolate genomes (n = 147). Genome pairs (species shown along the x axis) are grouped by genus. Empty green circles depict MAG estimates, black filled circles depict isolate genome estimates. See Methods for a full description of the functional categories and their genomic profiling.

**Supplementary Figure 3 – Pangenome characteristics of the ten species studied for convergent adaptation to industrialization.**

A. Count of the number of genes in the core-, accessory- and cloud-genome of the ten species of interest. See Methods for the definition and reconstruction of each pangenome category.
B. Count of the number of genes in pangenome categories as a function of the number of strain-level genome bins (StGBs) per species. Linear regressions per pangenome categories were calculated – core-genome: beta = -2.8, p-val = 0.8; accessory-genome: beta = 4.8, p-val = 0.9; cloud-genome: beta = 282, p-val = 5.59e-05. Size of core and accessory genomes are stable across species irrespective of the number of StGBs. The size of the cloud genome is positively correlated to the number of sampled StGBs.

**Supplementary Figure 4 – Gene tree-Species tree reconciliations reveal increasing gene content along lineages of industrialized hosts.**

The panel depicts the evolution of the number of genes along the phylogeny of BFRA, BUNI, PDIS, PMER and BWEX, based on the reconciliation-aware reconstruction of ancestral gene contents. Reconciliations were calculated from the set of gene families present in at least four StGBs (tips of the tree). Overall trends for an increase in gene content along lineages occurring in industrialized hosts is observed (Fig. 4), and is statistically significant for BFRA and PMER (Fig. 4).

**Supplementary Figure 5 – Gene enrichment analysis based on industrialization level for BTHE, BXYL, PDOR, PMER and BWEX.**

Gene enrichment analysis performed for BTHE, BXYL, PDOR, PMER and BWEX, based on categorical and continuous levels of industrialization. Data is presented as in Fig. 5. In brief, gene profiles were coded as presence/absence data and were correlated to host industrialization status, controlling for phylogeny. Significant hits (q-value < 0.05) are colored coded based on host lifestyle. Volcano plots showing data for all genes (top) and effect size plots of top hits (bottom) are shown. The number of statistically significant genes is shown next to each species acronym. Differentially enriched genes also correlated with PC1 Lifestyle (continuous proxy of industrialization) are shown in plain circles.

**Supplementary Figure 6 – Convergent elevation of gene-level Ka/Ks based on industrialization status.**

Evidence for convergent positive selection on individual genes based on host industrialization status. The panel shows a heatmap of Ka/Ks differences for 90% gene families between industrialized and non-industrialized strains across species (x-axis). Genes with higher Ka/Ks values in industrialized strains are shown in purple, and those with higher values in non-industrialized strains are shown in green. Absent genes shown in white. Ka/Ks differences are displayed on a square-root–transformed color gradient. Mean Ka/Ks differences across species are shown on the left of the heatmap. Gene family annotations (preferred gene names or COG/KO IDs) are displayed on the y-axis. Genes with consistent Delta Ka/Ks signs across 8 or 7 of 10 species are shown at the top and bottom, respectively.

**Supplementary Figure 7 – Convergent signals of SNV-host lifestyle associations across bacterial species at the level of KEGG KO categories.**

Convergent signals of SNV-host lifestyle associations across bacterial species at the level of KEGG KOs. Each tile represents a KEGG KO that contains at least one 90% gene family with host industrialization-associated SNVs. KEGG KOs are shown along the y-axis, and species along the x-axis, cells are color-coded by species identity. All SNV data can be found in Supp. Table 7.

### Supplementary Tables

- Supp. Table 1 – Metadata of isolate genomes and donor participants.
- Supp. Table 2 – Profiles and comparisons of genomic features between paired MAGs and isolate genomes.
- Supp. Table 3 – Pangenome fluidity estimates across ten species.
- Supp. Table 4 – Counts of Gene tree - Species tree reconciliation events (speciations, duplications, losses, transfers, presence, originations, copies and singletons) for the ten species.
- Supp. Table 5 – Statistics and results of gene enrichment analyses performed on 90% similarity gene families in the ten species.
- Supp. Table 6 – Gene-level Ka/Ks estimates of 90% similarity gene families for the ten species.
- Supp. Table 7 – Statistics and results of variant-level associations with host industrialization for BFRA, BOVA, BUNI, PDIS and PVUL.

#### Acknowledgments

This work was supported by grants from the Center for Microbiome Informatics and Therapeutics at MIT and the Rasmussen Family Foundation.

M.P, M.R and M.G. received support from the Deutsche Forschungsgemeinschaft (DFG - German Science Foundation) within the Collaborative Research Center (CRC) 1182 on “The Origin and Function of Metaorganisms” (Project-ID 261376515 – SFB 1182, project C5.1 to M.G., project C5.2 to M.P., project C5.3 to M.R.).

The study received infrastructure support from the DFG (German Science Foundation) within the Cluster of Excellence 2167 “PrecisionMedicine in Chronic Inflammation (PMI)” (EXC 2167-390884018).

M.G. received funding from the European Research Council (ERC) under the European Union’s Horizon 2020 research and innovation programme (CoG VESICULOME, Grant agreement No. 101126254).

M.G., M.P. and A.F. received funding from the DFG – Project number 426660215 with the RU 5042 miTarget on “The microbiome as a therapeutic target in inflammatory bowel disease”, subprojects TP01 “Targeting intestinal yeasts and pathogenic yeast-responsive CD4+ T cells in Crohn’s disease” and TP02 “Ecology and function of synthetic bacterial communities for the understanding and modulation of IBD-associated microbiomes”.

The project received funding from the European Union under the Horizon Europe grant agreement No. 101095470 (project miGut-Health). Views and opinions expressed are however those of the author(s) only and do not necessarily reflect those of the European Union nor European Health and Digital Executive Agency (HaDEA). Neither the European Union nor HaDEA can be held responsible for them.

G.J.Sz and L.L.Sz. are grateful for the help and support provided by the Scientific Computing and Data Analysis section of Core Facilities at OIST.

R. J. X. acknowledges funding from the National Institutes of Health (NIH) (Project Center for the Study of Inflammatory Bowel Disease at Massachusetts General Hospital - DK043351)

We thank Tamara Mason and the team at the Walkup Sequencing platform at the Broad Institute for support on sequencing efforts.

## Author contributions

Designed this study: M.P, M.G, M.R, E.J.A

Field administrative work & collection of data and samples: M.P, M.G, V.J, A.F, A.A, M.Y.A, S.O.A, Y.A.A, A.D, Y.A.N, F.I, Y.L.A.L, T.M.P, C.O, J.R, I.E.M

Performed processing of biospecimens and molecular work: M.P Biosample and Data curation: M.P, M.G, M.R, L.M, H.J, J.C

Data analysis: M.R, M.G, M.P, E.J.A, L.L.S, S.W, L.M-S, L.K.M, J.F.C, J.B, A.F, G.J.S

Supervision: M.G, M.P, E.J.A

Funding acquisition: M.P, M.G, E.J.A, R.J.X, A.F, J.B, G.J.S

Writing, original draft: M.G, M.P, M.R

Writing, review & editing: all authors

## References

1. Schnorr, S.L., Candela, M., Rampelli, S., Centanni, M., Consolandi, C., Basaglia, G., Turroni, S., Biagi, E., Peano, C., Severgnini, M., et al. (2014). Gut microbiome of the Hadza hunter-gatherers. Nat. Commun. 5, 3654. 10.1038/ncomms4654.

2. Carter, M.M., Olm, M.R., Merrill, B.D., Dahan, D., Tripathi, S., Spencer, S.P., Yu, F.B., Jain, S., Neff, N., Jha, A.R., et al. (2023). Ultra-deep sequencing of Hadza hunter-gatherers recovers vanishing gut microbes. Cell 186, 3111–3124.e13. 10.1016/j.cell.2023.05.046.

3. Smits, S.A., Leach, J., Sonnenburg, E.D., Gonzalez, C.G., Lichtman, J.S., Reid, G., Knight, R., Manjurano, A., Changalucha, J., Elias, J.E., et al. (2017). Seasonal cycling in the gut microbiome of the Hadza hunter-gatherers of Tanzania. Science 357, 802–806. 10.1126/science.aan4834.

4. Clemente, J.C., Pehrsson, E.C., Blaser, M.J., Sandhu, K., Gao, Z., Wang, B., Magris, M., Hidalgo, G., Contreras, M., Noya-Alarcón, Ó., et al. (2015). The microbiome of uncontacted Amerindians. Sci. Adv. 1, e1500183–e1500183. 10.1126/sciadv.1500183.

5. Maghini, D.G., Oduaran, O.H., Olubayo, L.A.I., Cook, J.A., Smyth, N., Mathema, T., Belger, C.W., Agongo, G., Boua, P.R., Choma, S.S.R., et al. (2025). Expanding the human gut microbiome atlas of Africa. Nature 638, 718–728. 10.1038/s41586-024-08485-8.

6. Poyet, M., Rühlemann, M., Schaan, A.P., Ma, Y., Moitinho-Silva, L., Wacker, E.M., Jebens, H., Patel, L., Nguyen, L.T.T., Zimmer, A., et al. (2025). Industrialization drives convergent microbial and physiological shifts in the human metaorganism. Co-submitted along this manuscript.

7. Bobay, L.-M., and Ochman, H. (2018). Factors driving effective population size and pan-genome evolution in bacteria. BMC Evol. Biol., 1–12. 10.1186/s12862-018-1272-4.

8. Roodgar, M., Good, B.H., Garud, N.R., Martis, S., Avula, M., Zhou, W., Lancaster, S.M., Lee, H., Babveyh, A., Nesamoney, S., et al. (2021). Longitudinal linked-read sequencing reveals ecological and evolutionary responses of a human gut microbiome during antibiotic treatment. Genome Res 31, 1433–1446. 10.1101/gr.265058.120.

9. Garud, N.R., Good, B.H., Hallatschek, O., and Pollard, K.S. (2019). Evolutionary dynamics of bacteria in the gut microbiome within and across hosts. Plos Biol 17, e3000102. 10.1371/journal.pbio.3000102.

10. Zhao, S., Lieberman, T.D., Poyet, M., Kauffman, K.M., Gibbons, S.M., Groussin, M., Xavier, R.J., and Alm, E.J. (2019). Adaptive Evolution within Gut Microbiomes of Healthy People. Cell Host Microbe 25, 656–667.e8. 10.1016/j.chom.2019.03.007.

11. Yaffe, E., and Relman, D.A. (2019). Tracking microbial evolution in the human gut using Hi-C reveals extensive horizontal gene transfer, persistence and adaptation. Nat. Microbiol. 16, 472–11. 10.1038/s41564-019-0625-0.

12. Poyet, M., Groussin, M., Gibbons, S.M., Avila-Pacheco, J., Jiang, X., Kearney, S.M., Perrotta, A.R., Berdy, B., Zhao, S., Lieberman, T.D., et al. (2019). A library of human gut bacterial isolates paired with longitudinal multiomics data enables mechanistic microbiome research. Nat. Med. 25, 1442– 1452. 10.1038/s41591-019-0559-3.

13. Browne, H.P., Forster, S.C., Anonye, B.O., Kumar, N., Neville, B.A., Stares, M.D., Goulding, D., and Lawley, T.D. (2016). Culturing of “unculturable” human microbiota reveals novel taxa and extensive sporulation. Nature 533, 543–546. 10.1038/nature17645.

14. Huang, Y., Sheth, R.U., Zhao, S., Cohen, L.A., Dabaghi, K., Moody, T., Sun, Y., Ricaurte, D., Richardson, M., Velez-Cortes, F., et al. (2023). High-throughput microbial culturomics using automation and machine learning. Nat. Biotechnol., 1–10. 10.1038/s41587-023-01674-2.

15. Forster, S.C., Kumar, N., Anonye, B.O., Almeida, A., Viciani, E., Stares, M.D., Dunn, M., Mkandawire, T.T., Zhu, A., Shao, Y., et al. (2019). A human gut bacterial genome and culture collection for improved metagenomic analyses. Nat. Biotechnol. 37, 186–192. 10.1038/s41587-018-0009-7.

16. Lin, X., Hu, T., Chen, J., Liang, H., Zhou, J., Wu, Z., Ye, C., Jin, X., Xu, X., Zhang, W., et al. (2023). The genomic landscape of reference genomes of cultivated human gut bacteria. Nat. Commun. 14, 1663. 10.1038/s41467-023-37396-x.

17. Goodman, A.L., Kallstrom, G., Faith, J.J., Reyes, A., Moore, A., Dantas, G., and Gordon, J.I. (2011). Extensive personal human gut microbiota culture collections characterized and manipulated in gnotobiotic mice. Proc Natl Acad Sci U A 108, 6252–6257. 10.1073/pnas.1102938108.

18. Zou, Y., Xue, W., Luo, G., Deng, Z., Qin, P., Guo, R., Sun, H., Xia, Y., Liang, S., Dai, Y., et al. (2019). 1,520 reference genomes from cultivated human gut bacteria enable functional microbiome analyses. Nat. Biotechnol., 1–15. 10.1038/s41587-018-0008-8.

19. Hitch, T.C.A., Masson, J.M., Pauvert, C., Bosch, J., Nüchtern, S., Treichel, N.S., Baloh, M., Razavi, S., Afrizal, A., Kousetzi, N., et al. (2025). HiBC: a publicly available collection of bacterial strains isolated from the human gut. Nat. Commun. 16, 4203. 10.1038/s41467-025-59229-9.

20. Huang, P., Dong, Q., Wang, Y., Tian, Y., Wang, S., Zhang, C., Yu, L., Tian, F., Gao, X., Guo, H., et al. (2024). Gut microbial genomes with paired isolates from China illustrate probiotic and cardiometabolic effects. Cell Genomics, 100559. 10.1016/j.xgen.2024.100559.

21. Groussin, M., Poyet, M., Sistiaga, A., Kearney, S.M., Moniz, K., Noel, M., Hooker, J., Gibbons, S.M., Segurel, L., Froment, A., et al. (2021). Elevated rates of horizontal gene transfer in the industrialized human microbiome. Cell 184, 2053–2067.e18. 10.1016/j.cell.2021.02.052.

22. Blanco-Míguez, A., Gálvez, E.J.C., Pasolli, E., De Filippis, F., Amend, L., Huang, K.D., Manghi, P., Lesker, T.-R., Riedel, T., Cova, L., et al. (2023). Extension of the Segatella copri complex to 13 species with distinct large extrachromosomal elements and associations with host conditions. Cell Host Microbe 31, 1804–1819.e9. 10.1016/j.chom.2023.09.013.

23. Pasolli, E., Asnicar, F., Manara, S., Zolfo, M., Karcher, N., Armanini, F., Beghini, F., Manghi, P., Tett, A., Ghensi, P., et al. (2019). Extensive Unexplored Human Microbiome Diversity Revealed by Over 150,000 Genomes from Metagenomes Spanning Age, Geography, and Lifestyle. Cell, 1–35. 10.1016/j.cell.2019.01.001.

24. Almeida, A., Nayfach, S., Boland, M., Strozzi, F., Beracochea, M., Shi, Z.J., Pollard, K.S., Sakharova, E., Parks, D.H., Hugenholtz, P., et al. (2020). A unified catalog of 204,938 reference genomes from the human gut microbiome. Nat. Biotechnol., 1–25. 10.1038/s41587-020-0603-3.

25. Nayfach, S., Shi, Z.J., Seshadri, R., Pollard, K.S., and Kyrpides, N.C. (2019). New insights from uncultivated genomes of the global human gut microbiome. Nature 568, 505–510. 10.1038/s41586-019-1058-x.

26. Karcher, N., Pasolli, E., Asnicar, F., Huang, K.D., Tett, A., Manara, S., Armanini, F., Bain, D., Duncan, S.H., Louis, P., et al. (2020). Analysis of 1321 Eubacterium rectale genomes from metagenomes uncovers complex phylogeographic population structure and subspecies functional adaptations. Genome Biol., 1–27. 10.1186/s13059-020-02042-y.

27. Shaiber, A., and Eren, A.M. (2019). Composite Metagenome-Assembled Genomes Reduce the Quality of Public Genome Repositories. mBio 10, 208–3. 10.1128/mBio.00725-19.

28. Meziti, A., Rodriguez-R, L.M., Hatt, J.K., Peña-Gonzalez, A., Levy, K., and Konstantinidis, K.T. (2021). The Reliability of Metagenome-Assembled Genomes (MAGs) in Representing Natural Populations: Insights from Comparing MAGs against Isolate Genomes Derived from the Same Fecal Sample. Appl. Environ. Microbiol. 87, e02593–20. 10.1128/AEM.02593-20.

29. Mageeney, C.M., Trubl, G., and Williams, K.P. (2022). Improved Mobilome Delineation in Fragmented Genomes. Front. Bioinforma. 2, 866850. 10.3389/fbinf.2022.866850.

30. Chen, L.-X., Anantharaman, K., Shaiber, A., Eren, A.M., and Banfield, J.F. (2020). Accurate and complete genomes from metagenomes. Genome Res. 30, 315–333. 10.1101/gr.258640.119.

31. Ramos-Barbero, M.D., Martin-Cuadrado, A.-B., Viver, T., Santos, F., Martinez-Garcia, M., and Antón, J. (2019). Recovering microbial genomes from metagenomes in hypersaline environments: The Good, the Bad and the Ugly. Syst. Appl. Microbiol. 42, 30–40. 10.1016/j.syapm.2018.11.001.

32. Obregon-Tito, A.J., Tito, R.Y., Metcalf, J., Sankaranarayanan, K., Clemente, J.C., Ursell, L.K., Zech Xu, Z., Van Treuren, W., Knight, R., Gaffney, P.M., et al. (2015). Subsistence strategies in traditional societies distinguish gut microbiomes. Nat. Commun. 6, 6505. 10.1038/ncomms7505.

33. Wibowo, M.C., Yang, Z., Borry, M., Hübner, A., Huang, K.D., Tierney, B.T., Zimmerman, S., Barajas-Olmos, F., Contreras-Cubas, C., García-Ortiz, H., et al. (2021). Reconstruction of ancient microbial genomes from the human gut. Nature 594, 234–239. 10.1038/s41586-021-03532-0.

34. Arif, S., Nirmalan, S., Alazizi, A., Mair-Meijers, H., Agyei, A., Afihene, M.Y., Asibey, S.O., Awuku, Y.A., Duah, A., Plymoth, A., et al. (2025). Host transcriptional responses to gut microbiome variation arising from urbanism. Co-submitted along this manuscript.

35. Looft, T., Levine, U.Y., and Stanton, T.B. (2013). Cloacibacillus porcorum sp. nov., a mucin-degrading bacterium from the swine intestinal tract and emended description of the genus Cloacibacillus. Int. J. Syst. Evol. Microbiol. 63, 1960–1966. 10.1099/ijs.0.044719-0.

36. Richardson, L., Allen, B., Baldi, G., Beracochea, M., Bileschi, M.L., Burdett, T., Burgin, J., Caballero-Pérez, J., Cochrane, G., Colwell, L.J., et al. (2023). MGnify: the microbiome sequence data analysis resource in 2023. Nucleic Acids Res. 51, D753–D759. 10.1093/nar/gkac1080.

37. Chaumeil, P.-A., Mussig, A.J., Hugenholtz, P., and Parks, D.H. (2020). GTDB-Tk: a toolkit to classify genomes with the Genome Taxonomy Database. Bioinformatics 36, 1925–1927. 10.1093/bioinformatics/btz848.

38. Starke, S., Harris, D.M.M., Paulay, A., Aden, K., and Waschina, S. (2025). Comparative analysis of amino acid auxotrophies and peptidase profiles in non-dysbiotic and dysbiotic small intestinal microbiomes. Comput. Struct. Biotechnol. J. 27, 821–831. 10.1016/j.csbj.2025.02.004.

39. Mise, K., and Iwasaki, W. (2022). Unexpected absence of ribosomal protein genes from metagenome-assembled genomes. ISME Commun. 2, 118. 10.1038/s43705-022-00204-6.

40. Rühlemann, M.C., Wacker, E.M., Ellinghaus, D., and Franke, A. (2022). MAGScoT : a fast, lightweight and accurate bin-refinement tool. Bioinformatics 38, 5430–5433. 10.1093/bioinformatics/btac694.

41. Smillie, C.S., Smith, M.B., Friedman, J., Cordero, O.X., David, L.A., and Alm, E.J. (2011). Ecology drives a global network of gene exchange connecting the human microbiome. Nature 480, 241–244. 10.1038/nature10571.

42. Chen, L., Zhao, N., Cao, J., Liu, X., Xu, J., Ma, Y., Yu, Y., Zhang, X., Zhang, W., Guan, X., et al. (2022). Short- and long-read metagenomics expand individualized structural variations in gut microbiomes. Nat. Commun. 13, 3175–12. 10.1038/s41467-022-30857-9.

43. Moss, E.L., Maghini, D.G., and Bhatt, A.S. (2020). Complete, closed bacterial genomes from microbiomes using nanopore sequencing. Nat. Biotechnol., 1–12. 10.1038/s41587-020-0422-6.

44. Morel, B., Williams, T.A., Stamatakis, A., and Szöllősi, G.J. (2024). AleRax: a tool for gene and species tree co-estimation and reconciliation under a probabilistic model of gene duplication, transfer, and loss. Bioinformatics 40, btae162. 10.1093/bioinformatics/btae162.

45. Mainprize, I.L., Bean, J.D., Bouwman, C., Kimber, M.S., and Whitfield, C. (2013). The UDP-glucose Dehydrogenase of Escherichia coli K-12 Displays Substrate Inhibition by NAD That Is Relieved by Nucleotide Triphosphates. J. Biol. Chem. 288, 23064–23074. 10.1074/jbc.M113.486613.

46. Lehrer, J., Vigeant, K.A., Tatar, L.D., and Valvano, M.A. (2007). Functional Characterization and Membrane Topology of *Escherichia coli* WecA, a Sugar-Phosphate Transferase Initiating the Biosynthesis of Enterobacterial Common Antigen and O-Antigen Lipopolysaccharide. J. Bacteriol. 189, 2618–2628. 10.1128/JB.01905-06.

47. Virolle, C., Goldlust, K., Djermoun, S., Bigot, S., and Lesterlin, C. (2020). Plasmid Transfer by Conjugation in Gram-Negative Bacteria: From the Cellular to the Community Level. Genes 11, 1239. 10.3390/genes11111239.

48. Wolff, R., and Garud, N.R. Pervasive selective sweeps across human gut microbiomes.

49. Groussin, M., Poyet, M., Sistiaga, A., Kearney, S.M., Moniz, K., Noel, M., Hooker, J., Gibbons, S.M., Ségurel, L., Froment, A., et al. (2021). Elevated rates of horizontal gene transfer in the industrialized human microbiome. Cell 184, 2053–2067.e18. 10.1016/j.cell.2021.02.052.

50. Haines-Menges, B., Whitaker, W.B., and Boyd, E.F. (2014). Alternative Sigma Factor RpoE Is Important for Vibrio parahaemolyticus Cell Envelope Stress Response and Intestinal Colonization. Infect. Immun. 82, 3667–3677. 10.1128/IAI.01854-14.

51. Gutierrez, J., Smith, R., and Pogliano, K. (2010). SpoIID-Mediated Peptidoglycan Degradation Is Required throughout Engulfment during *Bacillus subtilis* Sporulation. J. Bacteriol. 192, 3174–3186. 10.1128/JB.00127-10.

52. Heimesaat, M.M., Schmidt, A.-M., Mousavi, S., Escher, U., Tegtmeyer, N., Wessler, S., Gadermaier, G., Briza, P., Hofreuter, D., Bereswill, S., et al. (2020). Peptidase PepP is a novel virulence factor of *Campylobacter jejuni* contributing to murine campylobacteriosis. Gut Microbes 12, 1770017. 10.1080/19490976.2020.1770017.

53. Paterson, G.K., Cone, D.B., Northen, H., Peters, S.E., and Maskell, D.J. (2009). Deletion of the gene encoding the glycolytic enzyme triosephosphate isomerase (*tpi*) alters morphology of *Salmonella enterica* serovar Typhimurium and decreases fitness in mice. FEMS Microbiol. Lett. 294, 45–51. 10.1111/j.1574-6968.2009.01553.x.

54. Xia, Y., Wang, D., Pan, X., Xia, B., Weng, Y., Long, Y., Ren, H., Zhou, J., Jin, Y., Bai, F., et al. (2020). TpiA is a Key Metabolic Enzyme That Affects Virulence and Resistance to Aminoglycoside Antibiotics through CrcZ in Pseudomonas aeruginosa. mBio 11, e02079–19. 10.1128/mBio.02079-19.

55. Urbonavièius, J., Durand, J.M.B., and Björk, G.R. (2002). Three Modifications in the D and T Arms of tRNA Influence Translation in *Escherichia coli* and Expression of Virulence Genes in *Shigella flexneri*. J. Bacteriol. 184, 5348–5357. 10.1128/JB.184.19.5348-5357.2002.

56. Noel, H.R., Keerthi, S., Ren, X., Winkelman, J.D., Troutman, J.M., and Palmer, L.D. (2024). Genetic synergy between *Acinetobacter baumannii* undecaprenyl phosphate biosynthesis and the Mla system impacts cell envelope and antimicrobial resistance. mBio 15, e02804–23. 10.1128/mbio.02804-23.

57. Verstraete, M.M., Perez-Borrajero, C., Brown, K.L., Heinrichs, D.E., and Murphy, M.E.P. (2018). SbnI is a free serine kinase that generates -phospho-l-serine for staphyloferrin B biosynthesis in. J. Biol. Chem. 293, 6147–6160. 10.1074/jbc.RA118.001875.

58. Révora, V., Marchesini, M.I., and Comerci, D.J. (2020). *Brucella abortus* Depends on L-Serine Biosynthesis for Intracellular Proliferation. Infect. Immun. 88, e00840–19. 10.1128/IAI.00840-19.

59. Croucher, N.J., Page, A.J., Connor, T.R., Delaney, A.J., Keane, J.A., Bentley, S.D., Parkhill, J., and Harris, S.R. (2015). Rapid phylogenetic analysis of large samples of recombinant bacterial whole genome sequences using Gubbins. Nucleic Acids Res 43, e15–e15. 10.1093/nar/gku1196.

60. Sonnenburg, J.L., and Sonnenburg, E.D. (2019). Vulnerability of the industrialized microbiota. Science 366, eaaw9255. 10.1126/science.aaw9255.

61. Vatanen, T., Kostic, A.D., d’Hennezel, E., Siljander, H., Franzosa, E.A., Yassour, M., Kolde, R., Vlamakis, H., Arthur, T.D., Hämäläinen, A.-M., et al. (2016). Variation in Microbiome LPS Immunogenicity Contributes to Autoimmunity in Humans. Cell 165, 842–853. 10.1016/j.cell.2016.04.007.

62. Martin, M. (2011). Cutadapt removes adapter sequences from high-throughput sequencing reads. EMBnet J, 10–12.

63. Bolger, A.M., Lohse, M., and Usadel, B. (2014). Trimmomatic: a flexible trimmer for Illumina sequence data. Bioinformatics 30, 2114–2120. 10.1093/bioinformatics/btu170.

64. Bankevich, A., Nurk, S., Antipov, D., Gurevich, A.A., Dvorkin, M., Kulikov, A.S., Lesin, V.M., Nikolenko, S.I., Pham, S., Prjibelski, A.D., et al. (2012). SPAdes: a new genome assembly algorithm and its applications to single-cell sequencing. J Comput Biol 19, 455–477. 10.1089/cmb.2012.0021.

65. Boetzer, M., Henkel, C.V., Jansen, H.J., Butler, D., and Pirovano, W. (2011). Scaffolding pre-assembled contigs using SSPACE. Bioinformatics 27, 578–579. 10.1093/bioinformatics/btq683.

66. Nadalin, F., Vezzi, F., and Policriti, A. (2012). GapFiller: a de novo assembly approach to fill the gap within paired reads. BMC Bioinformatics 13, S8. 10.1186/1471-2105-13-S14-S8.

67. Li, D., Liu, C.-M., Luo, R., Sadakane, K., and Lam, T.-W. (2015). MEGAHIT: an ultra-fast single-node solution for large and complex metagenomics assembly via succinct *de Bruijn* graph. Bioinformatics 31, 1674–1676. 10.1093/bioinformatics/btv033.

68. Wu, Y.-W., Simmons, B.A., and Singer, S.W. (2016). MaxBin 2.0: an automated binning algorithm to recover genomes from multiple metagenomic datasets. Bioinformatics 32, 605–607. 10.1093/bioinformatics/btv638.

69. Kang, D.D., Li, F., Kirton, E., Thomas, A., Egan, R., An, H., and Wang, Z. (2019). MetaBAT 2: an adaptive binning algorithm for robust and efficient genome reconstruction from metagenome assemblies. PeerJ 7, e7359. 10.7717/peerj.7359.

70. Alneberg, J., Bjarnason, B.S., De Bruijn, I., Schirmer, M., Quick, J., Ijaz, U.Z., Lahti, L., Loman, N.J., Andersson, A.F., and Quince, C. (2014). Binning metagenomic contigs by coverage and composition. Nat. Methods 11, 1144–1146. 10.1038/nmeth.3103.

71. Nissen, J.N., Johansen, J., Allesøe, R.L., Sønderby, C.K., Armenteros, J.J.A., Grønbech, C.H., Jensen, L.J., Nielsen, H.B., Petersen, T.N., Winther, O., et al. (2021). Improved metagenome binning and assembly using deep variational autoencoders. Nat. Biotechnol. 39, 555–560. 10.1038/s41587-020-00777-4.

72. Parks, D.H., Imelfort, M., Skennerton, C.T., Hugenholtz, P., and Tyson, G.W. (2015). CheckM: assessing the quality of microbial genomes recovered from isolates, single cells, and metagenomes. Genome Res 25, 1043–1055. 10.1101/gr.186072.114.

73. Srivastava, A., Malik, L., Smith, T., Sudbery, I., and Patro, R. (2019). Alevin efficiently estimates accurate gene abundances from dscRNA-seq data. Genome Biol. 20, 65. 10.1186/s13059-019-1670-y.

74. Zimmermann, J., Kaleta, C., and Waschina, S. (2021). gapseq: informed prediction of bacterial metabolic pathways and reconstruction of accurate metabolic models. Genome Biol 22, 81–35. 10.1186/s13059-021-02295-1.

75. Starke, S., Harris, D.M.M., Zimmermann, J., Schuchardt, S., Oumari, M., Frank, D., Bang, C., Rosenstiel, P., Schreiber, S., Frey, N., et al. (2023). Amino acid auxotrophies in human gut bacteria are linked to higher microbiome diversity and long-term stability. ISME J. 17, 2370–2380. 10.1038/s41396-023-01537-3.

76. Feldgarden, M., Brover, V., Gonzalez-Escalona, N., Frye, J.G., Haendiges, J., Haft, D.H., Hoffmann, M., Pettengill, J.B., Prasad, A.B., Tillman, G.E., et al. (2021). AMRFinderPlus and the Reference Gene Catalog facilitate examination of the genomic links among antimicrobial resistance, stress response, and virulence. Sci. Rep. 11, 12728. 10.1038/s41598-021-91456-0.

77. Liu, B., Zheng, D., Jin, Q., Chen, L., and Yang, J. (2019). VFDB 2019: a comparative pathogenomic platform with an interactive web interface. Nucleic Acids Res. 47, D687–D692. 10.1093/nar/gky1080.

78. Zheng, J., Ge, Q., Yan, Y., Zhang, X., Huang, L., and Yin, Y. (2023). dbCAN3: automated carbohydrate-active enzyme and substrate annotation. Nucleic Acids Res. 51, W115–W121. 10.1093/nar/gkad328.

79. Abby, S.S., Denise, R., and Rocha, E.P.C. (2024). Identification of Protein Secretion Systems in Bacterial Genomes Using MacSyFinder Version 2. In Bacterial Secretion Systems Methods in Molecular Biology., L. Journet and E. Cascales, eds. (Springer US), pp. 1–25. 10.1007/978-1-0716-3445-5_1.

80. Xie, Z., and Tang, H. (2017). ISEScan: automated identification of insertion sequence elements in prokaryotic genomes. Bioinformatics 33, 3340–3347. 10.1093/bioinformatics/btx433.

81. Camargo, A.P., Roux, S., Schulz, F., Babinski, M., Xu, Y., Hu, B., Chain, P.S.G., Nayfach, S., and Kyrpides, N.C. (2023). Identification of mobile genetic elements with geNomad. Nat. Biotechnol. 10.1038/s41587-023-01953-y.

82. Camarillo-Guerrero, L.F., Almeida, A., Rangel-Pineros, G., Finn, R.D., and Lawley, T.D. (2021). Massive expansion of human gut bacteriophage diversity. Cell 184, 1098–1109.e9. 10.1016/j.cell.2021.01.029.

83. Nayfach, S., Camargo, A.P., Schulz, F., Eloe-Fadrosh, E., Roux, S., and Kyrpides, N.C. (2021). CheckV assesses the quality and completeness of metagenome-assembled viral genomes. Nat. Biotechnol. 39, 578–585. 10.1038/s41587-020-00774-7.

84. Preska Steinberg, A., and Kussell, E. (2025). How recombination and clonal evolution shape bacterial lineages and genomes. GENETICS, iyaf115. 10.1093/genetics/iyaf115.

85. Cingolani, P., Platts, A., Wang, L.L., Coon, M., Nguyen, T., Wang, L., Land, S.J., Lu, X., and Ruden, D.M. (2012). A program for annotating and predicting the effects of single nucleotide polymorphisms, SnpEff: SNPs in the genome of Drosophila melanogaster strain w1118; iso-2; iso-3. Fly (Austin) 6, 80–92. 10.4161/fly.19695.

86. Minh, B.Q., Schmidt, H.A., Chernomor, O., Schrempf, D., Woodhams, M.D., Von Haeseler, A., and Lanfear, R. (2020). IQ-TREE 2: New Models and Efficient Methods for Phylogenetic Inference in the Genomic Era. Mol. Biol. Evol. 37, 1530–1534. 10.1093/molbev/msaa015.

87. Seemann, T. (2014). Prokka: rapid prokaryotic genome annotation. Bioinformatics 30, 2068–2069. 10.1093/bioinformatics/btu153.

88. Steinegger, M., and Söding, J. (2017). MMseqs2 enables sensitive protein sequence searching for the analysis of massive data sets. Nat. Biotechnol. 35, 1026–1028. 10.1038/nbt.3988.

89. Cantalapiedra, C.P., Hernández-Plaza, A., Letunic, I., Bork, P., and Huerta-Cepas, J. (2021). eggNOG-mapper v2: Functional Annotation, Orthology Assignments, and Domain Prediction at the Metagenomic Scale. Mol. Biol. Evol. 38, 5825–5829. 10.1093/molbev/msab293.

90. Katoh, K., and Standley, D.M. (2013). MAFFT Multiple Sequence Alignment Software Version 7: Improvements in Performance and Usability. Mol. Biol. Evol. 30, 772–780. 10.1093/molbev/mst010.

91. Kislyuk, A.O., Haegeman, B., Bergman, N.H., and Weitz, J.S. (2011). Genomic fluidity: an integrative view of gene diversity within microbial populations. BMC Genomics 12, 32. 10.1186/1471-2164-12-32.

92. Dewar, A.E., Hao, C., Belcher, L.J., Ghoul, M., and West, S.A. (2024). Bacterial lifestyle shapes pangenomes. Proc. Natl. Acad. Sci. 121, e2320170121. 10.1073/pnas.2320170121.

93. Szöllõsi, G.J., Rosikiewicz, W., Boussau, B., Tannier, E., and Daubin, V. (2013). Efficient exploration of the space of reconciled gene trees. Syst Biol 62, 901–912. 10.1093/sysbio/syt054.

94. Huerta-Cepas, J., Serra, F., and Bork, P. (2016). ETE 3: Reconstruction, Analysis, and Visualization of Phylogenomic Data. Mol. Biol. Evol. 33, 1635–1638. 10.1093/molbev/msw046.

95. Hoang, D.T., Chernomor, O., Von Haeseler, A., Minh, B.Q., and Vinh, L.S. (2018). UFBoot2: Improving the Ultrafast Bootstrap Approximation. Mol. Biol. Evol. 35, 518–522. 10.1093/molbev/msx281.

96. Kluge, A.G., and Farris, J.S. (1969). Quantitative Phyletics and the Evolution of Anurans. Syst. Biol. 18, 1–32. 10.1093/sysbio/18.1.1.

97. Tung Ho, L.S., and Ané, C. (2014). A Linear-Time Algorithm for Gaussian and Non-Gaussian Trait Evolution Models. Syst. Biol. 63, 397–408. 10.1093/sysbio/syu005.

